# Symmetric Dimethylarginine Modification Drives Shared and Divergent Antigen Reactivity in Lupus

**DOI:** 10.64898/2026.09.22.753525

**Authors:** Pan Liu, Huixian Li, Andrew Zu-Sern Wei, Jin G. Park, Wanhong Lu, Xinfang Xie, Jing Jin

## Abstract

Systemic lupus erythematosus (SLE) features a broad array of autoantibodies against nuclear components, but the mechanisms driving this diversity are poorly understood. We investigated whether posttranslational arginine methylation creates common neo epitopes that could unify this response. Using methylome wide peptide arrays, we identified anti symmetric dimethylarginine (SDMA) autoantibodies in 30-40% of SLE patients, with individualized reactivity fingerprints. A single anti SDMA monoclonal antibody from MRL lpr mice recognized multiple SDMA modified antigens, and EBNA 1, an Epstein Barr virus protein, bound SLE plasma in an SDMA dependent manner, pointing to potential molecular mimicry. In a validation cohort of 201 patients, anti SDMA titers correlated with SLEDAI 2K scores and inversely with C3/C4; seropositivity was overrepresented in younger patients, those with renal involvement, and anti Sm-positive individuals. These results establish that anti SDMA responses are common in SLE, target methylated arginine motifs across self and viral proteins, and serve as markers of active, complement consuming disease. The patient specific recognition of SDMA epitopes may underlie the serological heterogeneity that defines SLE.

## Introduction

Systemic lupus erythematosus (SLE) autoimmunity is frequently associated with antinuclear antibodies (ANAs) targeting a diverse array of nuclear antigens, including dsDNA, histones, centromere proteins, topoisomerase I, and numerous RNA-binding proteins (RBPs) such as Sm, RNP, SSA/Ro, SSB/La, and ribosomal proteins(1, 2). Specific correlations exist between certain autoantibodies and organ involvement(3-5). For instance, anti-dsDNA antibodies are commonly linked to renal involvement(6), anti-Sm antibodies are highly specific for SLE(7), whereas anti-U1 snRNP antibodies are found in both SLE and mixed connective tissue disease (MCTD)(8), and anti-SSA/Ro and anti-SSB/La antibodies are typically associated with subacute cutaneous lupus and Sjögren disease (SjD)(9).

Despite the vast diversity of RBPs, they form nuclear structures that are often distinguishable in ANA tests. SSA/Ro and SSB/La are predominantly localized to the nucleolus(10), whereas RNP—referring to a highly diverse group of RNA granules—encompasses a broad range of ribonucleoprotein particles(11). The Sm core is a seven-protein complex (subunits B/BM, D1-D3, E, F, and G) essential for the biogenesis of the U small nuclear ribonucleoprotein (U-snRNP) spliceosome, a complex containing at least 50 distinct proteins(12). Although the antigenic nature of these RNPs in lupus remains unclear, many autoimmune diseases, including MCTD, SS, rheumatoid arthritis (RA), systemic sclerosis (SSc), and dermatomyositis (DM), share overlapping serological patterns with SLE(13, 14). Such targets include SmD1, SmD3, nucleolin (an hnRNP), and the EBV antigen EBNA 1, a well established SLE autoantigen. Notably, these antigens share common post translational arginine methylation, which has been linked to the autoantibody response(15-17). In 2000, Brahms and colleagues reported that anti-Sm autoantibodies recognize symmetrical dimethylarginines (SDMAs) within arginine-glycine-rich sequences on SmD1 and SmD3(18). Subsequently, anti-Sm-positive SLE sera were found to recognize hnRNP D-like protein and cellular nucleic acid binding protein (CNBP) through similar methylarginine epitopes(19). Moreover, van Vliet et al. demonstrated that anti-Sm IgG cross-reacts with Epstein-Barr nuclear antigen 1 (EBNA1) via shared SDMA epitopes, suggesting a broad anti-posttranslational modification response in SLE(15).

From a broader perspective, this study was designed to address three fundamental questions: (1) whether shared epitope features, particularly posttranslational modifications such as arginine methylation, account for the large number and variety of SLE autoantigens; (2) whether the heterogeneity of autoantigen reactivity among SLE patients arises from stochastic recognition of flanking sequences surrounding a conserved methylarginine residue; and (3) whether SLE can be conceptualized as a clonal disease, wherein a single clonal autoantibody population broadly cross-reacts with multiple antigens through recognition of a shared methylarginine epitope.

## Results

### Heterogeneity in Autoantibody Specificity Among SLE Patients

We recruited a discovery cohort of 31 SLE patients and investigated their cellular antigen patterns using immunoblotting (IB) and immunofluorescence (IF) staining of HEp-2 cells with plasma samples (Figure 1, and patient information is provided in Table 1). Immunoblotting against cell lysates revealed a wide variety of reactive band patterns in a majority of patients (23/31), with distinct differences in band intensities and molecular weights (Figure 1A). In contrast, two healthy control plasma pools showed no reactivity to HEp-2 cell lysate. Immunofluorescence staining detected antinuclear antibody reactivity in 21 of the 31 patients (Figure 1B, Supplemental Figure S1 and S2). As expected, nuclear staining patterns were diverse and could be broadly categorized as homogeneous (diffuse), speckled, nucleolar, peripheral (rim), or combinations thereof.

**Figure 1.**
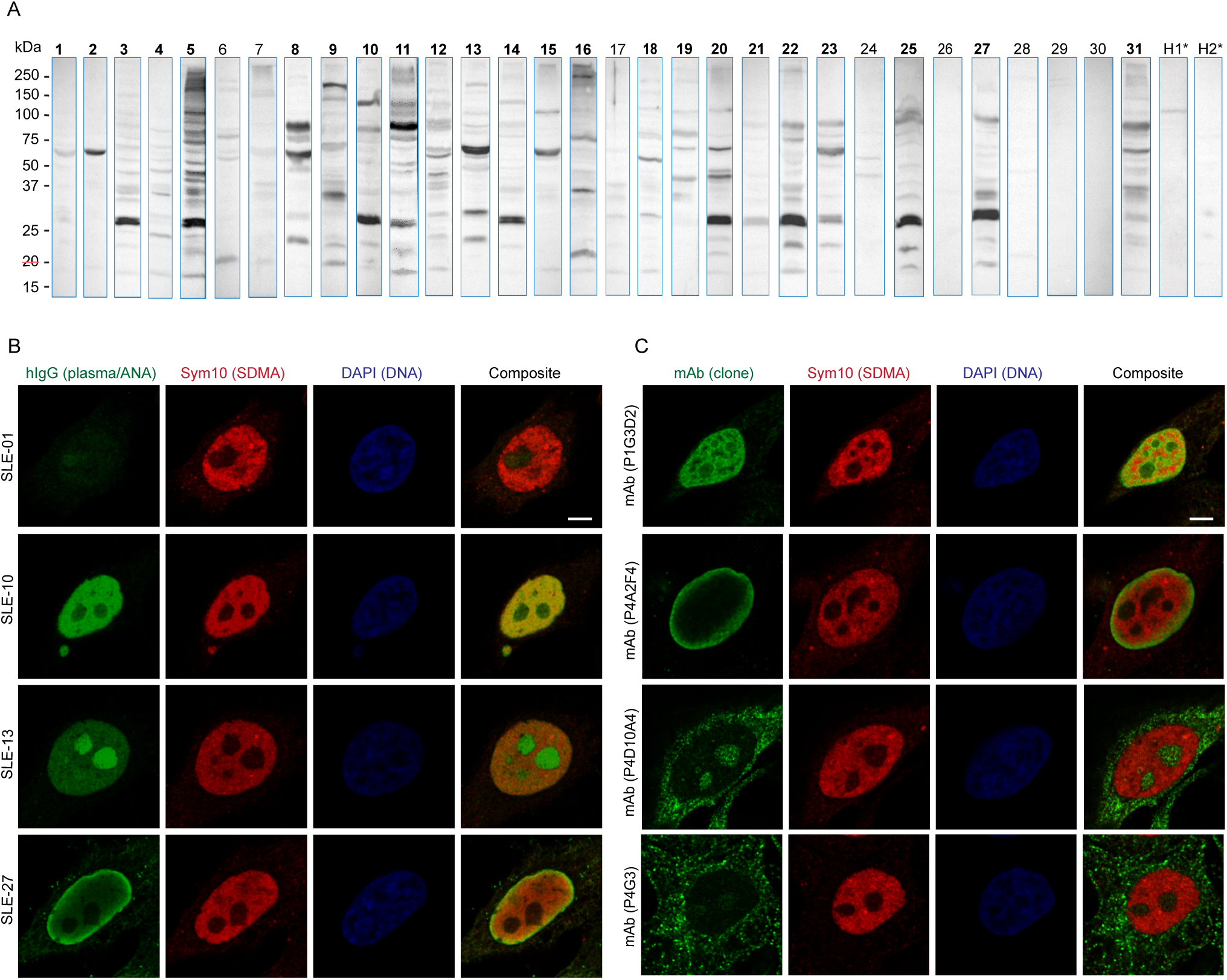
Diverse autoantigen patterns in patients with systemic lupus erythematosus. Plasma samples from 31 SLE patients and two healthy controls were analyzed using two complementary approaches: (A) immunoblotting of HEp 2 cell lysates separated by SDS PAGE, and (B) indirect immunofluorescence on fixed HEp 2 cells (ANA test; additional images in Supplemental Figure S1 and S2). (A) Immunoblotting revealed distinct antibody-reactive band patterns across patients, with variable band intensities. Several patients (e.g., SLE 01, 06, 07, 17, 24, 28, 29, 30) showed few bands, similar to healthy controls. (B) In the ANA test, selected plasma samples (SLE 01, 10, 13, 27) displayed contrasting nuclear staining patterns (green). A commercial monoclonal anti SDMA antibody (SYM10) was used as a counterstain in a separate channel (red). (C) Four monoclonal antibodies (mAbs) cloned from MRL lpr mice were used to stain HEp-2 cells. These mAbs exhibited distinct staining patterns, some of which resembled those observed with SLE patient plasma. For example, mAb P 1G3D2 resembled SLE 10, and mAb P4A2F4 resembled SLE 27.

**Table 1:** Baseline Characteristics and Anti-SDMA Autoantibody Findings in the Discovery Cohort (n=31)

| Pt<br>(n=31) | Sex | Age | Race | Disease<br>duration | Anti-SDMA | Anti-dsDNA | Anti-Sm<br>antigen | Anti-EBV | C3* | C4* | eGFR* |
| --- | --- | --- | --- | --- | --- | --- | --- | --- | --- | --- | --- |
| SLE-01 | M | 43 | White | 26y | - | - | - | + | 0.85 | 0.13 | 38 |
| SLE-02 | M | 60 | Other | NA | - | - | - | + | 0.99 | 0.40 | 44 |
| SLE-03 | F | 32 | Black | 18y | + | + | + | + | 0.52 | 0.08 | 98 |
| SLE-04 | F | 31 | Black | NA | - | + | - | + | NA | NA | 111 |
| SLE-05 | F | 63 | Other | >20y | + | + | + | + | 0.72 | 0.04 | 48 |
| SLE-06 | F | 59 | White | 20y | - | - | - | + | 0.97 | 0.23 | 30 |
| SLE-07 | F | 55 | Asian | 25y | + | + | - | + | 1.08 | 0.26 | 44 |
| SLE-08 | F | 40 | Black | NA | - | - | - | + | 0.40 | 0.03 | 87 |
| SLE-09 | F | 26 | Asian | 4y | + | + | - | + | 0.32 | 0.06 | 81 |
| SLE-10 | F | 23 | Black | 3y | + | - | + | + | 0.96 | 0.25 | 69 |
| SLE-11 | F | 42 | Other | 19y | + | - | + | + | 1.14 | 0.12 | 78 |
| SLE-12 | F | 33 | Other | 23y | + | +/- | - | + | 1.23 | 0.25 | 14 |
| SLE-13 | F | 58 | Black | 14y | + | - | - | + | 1.05 | 0.28 | 30 |
| SLE-14 | F | 35 | Other | 6y | - | +/- | + | + | 1.04 | 0.19 | 58 |
| SLE-15 | F | 45 | Black | 9y | - | +/- | - | + | 1.14 | 0.38 | 85 |
| SLE-16 | F | 39 | Black | 6y | - | + | - | + | 0.51 | 0.15 | 15 |
| SLE-17 | F | 66 | Black | 28y | - | + | - | + | 1.46 | 0.20 | 39 |
| SLE-18 | F | 37 | White | 10y | - | - | - | + | 1.00 | 0.30 | 67 |
| SLE-19 | F | 56 | Black | 27y | - | + | + | + | 1.42 | 0.22 | 98 |
| SLE-20 | F | 60 | Black | 9y | + | - | + | + | 0.73 | 0.21 | 26 |
| SLE-21 | F | 27 | Hispanic | 0y | - | + | - | + | 0.55 | 0.09 | 109 |
| SLE-22 | F | 50 | Black | 6y | + | + | + | + | 1.19 | 0.23 | 75 |
| SLE-23 | F | 26 | White | NA | + | + | + | + | 0.38 | 0.08 | 51 |
| SLE-24 | F | 53 | Other | 19y | - | - | + | + | 1.24 | 0.26 | 7 |
| SLE-25 | F | 27 | White | 10y | + | - | + | + | 0.89 | 0.07 | 76 |
| SLE-26 | F | 64 | White | NA | - | - | - | + | NA | NA | 16 |
| SLE-27 | F | 32 | Black | 1y | + | + | + | + | 0.72 | 0.13 | 110 |
| SLE-28 | F | 25 | Hispanic | 4y | - | +/- | - | + | 0.85 | 0.15 | 16 |
| SLE-29 | F | 54 | Asian | 21y | - | - | - | + | 0.92 | 0.14 | 64 |
| SLE-30 | M | 29 | White | 2y | - | +/- | - | - | 0.82 | 0.19 | 70 |
| SLE-31 | F | 42 | Black | 13y | - | + | + | + | 1.22 | 0.24 | 64 |

\*Reference range: C3 (75-175 g/L), C4 (14-40 g/L), eGFR ( $\geq 90$ mL/min/1.73 m<sup>2</sup>)

### Arginine-Glycine Dipeptide Repeats Are Frequent in RNA-Binding Proteins and, Upon Methylation, Serve as Neoantigens Targeted by Lupus Autoantibodies

As we focused on nuclear RBPs to identify shared sequence features, it became apparent that despite often comprising structurally unrelated domains, these proteins frequently harbor unstructured regions populated with arginine-glycine (RG) dipeptide repeats (Figure 2A). To further characterize the correlation between RG repeats and RNA-binding proteins, we searched the genome for proteins containing characteristic RG-repeat motifs. Gene Ontology (GO) analysis of the resulting list revealed an enrichment of RG repeats in nuclear proteins involved in RNA biogenesis (Supplemental Figure S3), which coincides with ANA-associated structures such as RNP granules, nuclear bodies, nuclear speckles, and nucleolus.

**Figure 2.**
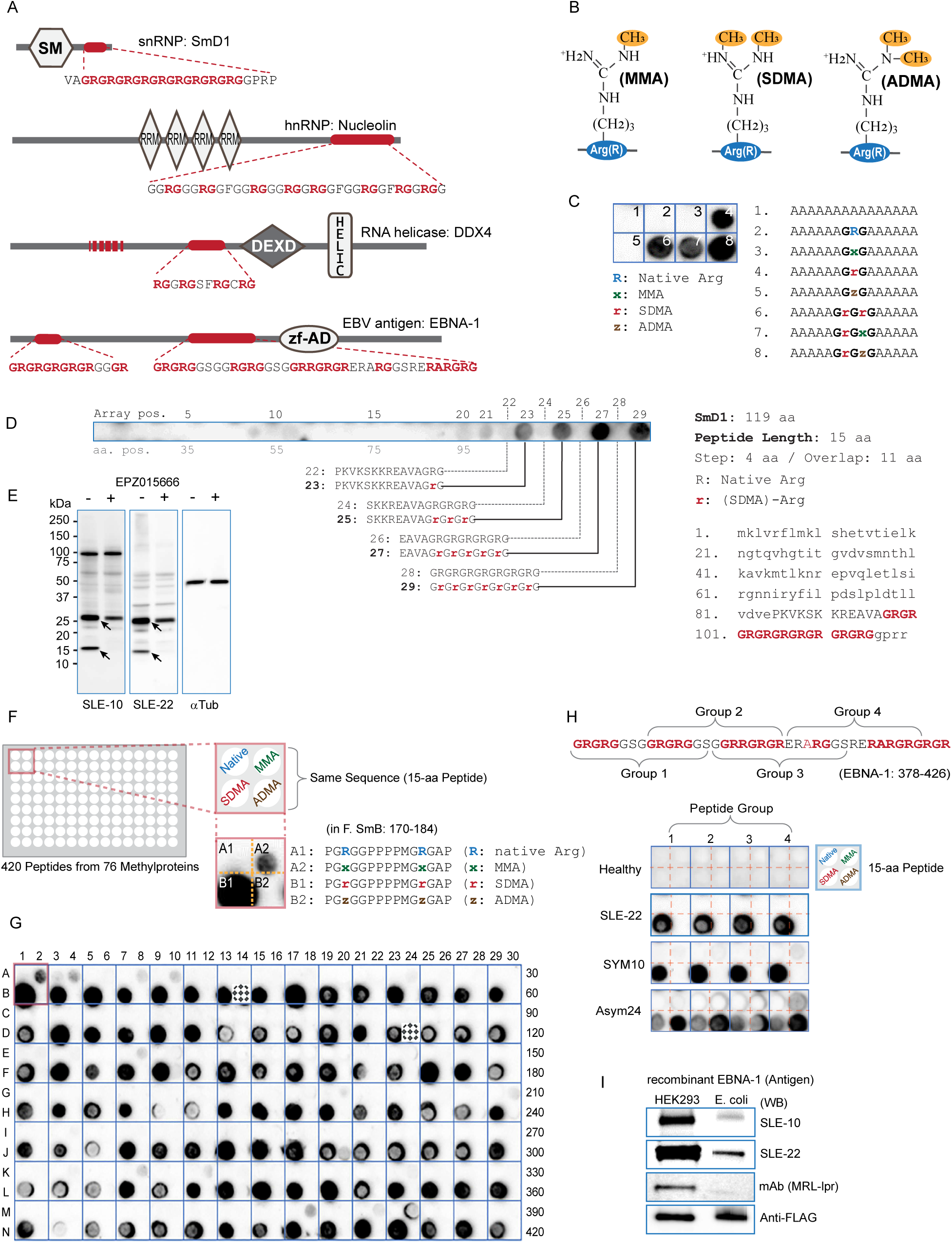
Symmetric dimethylarginine (SDMA) within Arg-Gly-rich sequences constitutes a reactive epitope for SLE autoantibodies. (A) Domain architectures of four representative RNA-binding proteins, each containing Arg-Gly-rich segments (red) of varying length and sequence. (B) Chemical structures of monomethylarginine (MMA), symmetric dimethylarginine (SDMA), and asymmetric dimethylarginine (ADMA), with methyl groups highlighted. (C) Eight synthetic peptides bearing different methylated arginine forms (or combinations thereof) were probed with plasma from SLE-22 and visualized with HRP-conjugated goat anti-human IgG (left panel). Peptides 4, 6, 7, and 8, all contained SDMA, reacted with SLE autoantibodies, whereas native, MMA-, and ADMA-containing peptides did not. (D) A tiling array of 29 peptides derived from human SmD1 (the RG-repeat region is highlighted, lower right). Peptides were designed with an 11-amino acid overlap (top right). Peptides 22-29, spanning the RG-repeats, were spotted in duplicate in either native or SDMA form (bottom). Only SDMA-containing peptides (spots 23, 25, 27, and 29) reacted with SLE-22 autoantibodies. (E) PRMT5 inhibitor EPZ015666 reduces specific immunoblot band intensities recognized by SLE-10 and SLE-22 plasma in HEp-2 cells. Arrows indicate bands with reduced intensity following inhibitor pretreatment. α-Tubulin (αTub) serves as a loading control. (F) Schematic of an array containing 105 peptides from 76 proteins identified in a human methylarginine proteome screen(35). Each peptide was synthesized in native, MMA, SDMA, or ADMA form. Representative data for SmB (amino acids 170-184) show strong reactivity of the SDMA-modified B1 peptide with SLE-22, weak reactivity for the MMA-modified A2 peptide, and no reactivity for native or ADMA forms. (G) Overview of the full array (420 peptides). SLE-22 autoantibodies reacted broadly with predominantly SDMA-containing peptides, with variable signal intensities. Checkered patches indicate two mistakenly synthesized sequences that were still included in the screen. (H) EBV antigen EBNA-1 contains two Arg-Gly-rich segments (as described in A). A 16-peptide array (divided into 4 groups) covering the longer segment (amino acids 378-426) was probed with healthy control plasma, SLE-22 plasma, or control antibodies SYM10 and ASM24. SLE-22 reacted with all four SDMA-containing peptides, mirroring the reactivity pattern of SYM10. (I) Full-length FLAG-tagged recombinant EBNA-1 was expressed in HEK293 cells (human cells that naturally produce arginine methylation, though with unknown frequency) and E. coli (prokaryotes do not generate arginine side-chain methylation). As expected, HEK293-derived EBNA-1 showed stronger reactivity with plasma from SLE-10 and SLE-22, as well as with a monoclonal antibody from the lupus-prone MRL-lpr mouse strain.

One key function of RG repeats is to mediate phase transitions of the constituent proteins(20-23), a process regulated in part by posttranslational methylation of arginine side chains(24-26). Arginine methylation is catalyzed by the protein arginine N-methyltransferase family (PRMT1-11 in humans)(27) and occurs in three forms: monomethylarginine (MMA), symmetric dimethylarginine (SDMA), and asymmetric dimethylarginine (ADMA) (Figure 2B)(28). Previous studies have shown that anti-Sm patient sera recognize these modified sequences(18, 19, 29). Examples of phase transitions in cells include the transient formation of stress granules containing translation initiation complexes and mRNAs, as well as nuclear organelles with key functions in RNA biogenesis(30-33).

We sought to experimentally characterize the role of arginine methylation in SLE antigenicity. To this end, we synthesized a set of eight peptides centered around a G-R-G motif, with the arginine residue presented either in its unmodified form or as MMA, SDMA, or ADMA. These peptides, containing one or two GRG motifs flanked by poly-alanine sequences, were spotted onto a membrane array. The membrane was then probed with plasma samples from SLE patients. While some samples showed no reactivity (data not shown), others exhibited antibody binding to specific modified peptides. For example, plasma from patient SLE-22 reacted only to peptides containing one or more SDMA residues (Figure 2C), indicating SDMA-dependent antibody recognition.

To further demonstrate that SLE plasma recognizes SDMA-modified epitopes rather than native autoantigens, we synthesized an additional set of 29 peptides, each 15 amino acids in length, based on the N- to C-terminal sequence of SmD1, a well-characterized lupus autoantigen(34). Notably, nine tandem RG repeats are located near the C-terminus of the sequence (Figure 2A, 2D). For each corresponding peptide, the arginine residues were presented either in their native form or as SDMA. The peptide set was probed with SLE-22 plasma, and lupus antibodies were clearly directed specifically toward SDMA-modified peptides containing the RG repeats (Figure 2D). These findings indicate that lupus antibodies target SmD1 exclusively through their methylated C-terminus rather than the native antigen. Consistent with this conclusion, pretreatment of HEp-2 cells with PRMT5 inhibitor EPZ015666 reduced the intensity of specific SLE-reactive bands (Figure 2E), further supporting that autoantibodies mainly recognize methylarginine-containing epitopes, in agreement with previous reports(15, 18).

### Global Anti-SDMA Reactivity Reveals Diverse Patterns Among SLE Patients

We next expanded our analysis of anti-methylarginine autoantibodies to a broader set of autoantigens previously identified by mass spectrometry as containing methylated arginine residues(35). A total of 78 proteins, represented by 105 peptides encompassing those methylarginine sites (Supplemental Table S1), were included. Each peptide sequence was synthesized on an array (Figure 2F), with all arginine residues presented either unmodified or modified as MMA, SDMA, or ADMA. The arrays were then probed with plasma samples from SLE patients. Overall, antibodies predominantly recognized SDMA-containing peptides (Figure 2G shows an example from patient SLE-22), highlighting the dominant role of SDMA in the SLE autoantibody response, an observation that extends beyond established SLE autoantigens.

In the next array, we included only the SDMA-modified peptides alongside their corresponding native sequences from the same set (Supplemental Table S1). To validate array quality, we performed a series of tests using a commercial rabbit polyclonal methylarginine antibody (SYM10), a mouse monoclonal antibody against SNRPB/SmB (Y12), and a commercial anti-SmD1 antibody (Figure 3A). As expected, polyclonal SYM10 broadly recognized all 105 SDMA-modified peptides but only 2-3 native peptides, likely due to incidental cross-reactivity. Monoclonal Y12(36) recognized a subset of SDMA peptides with a wide range of signal intensities, likely reflecting its selectivity for flanking sequences that, together with SDMA, form the antigenic epitope. In contrast, the commercial anti-SmD1 antibody recognized only a small subset of native sequences, potentially due to the presence of RG repeats resembling those found in SmD1.

**Figure 3.**
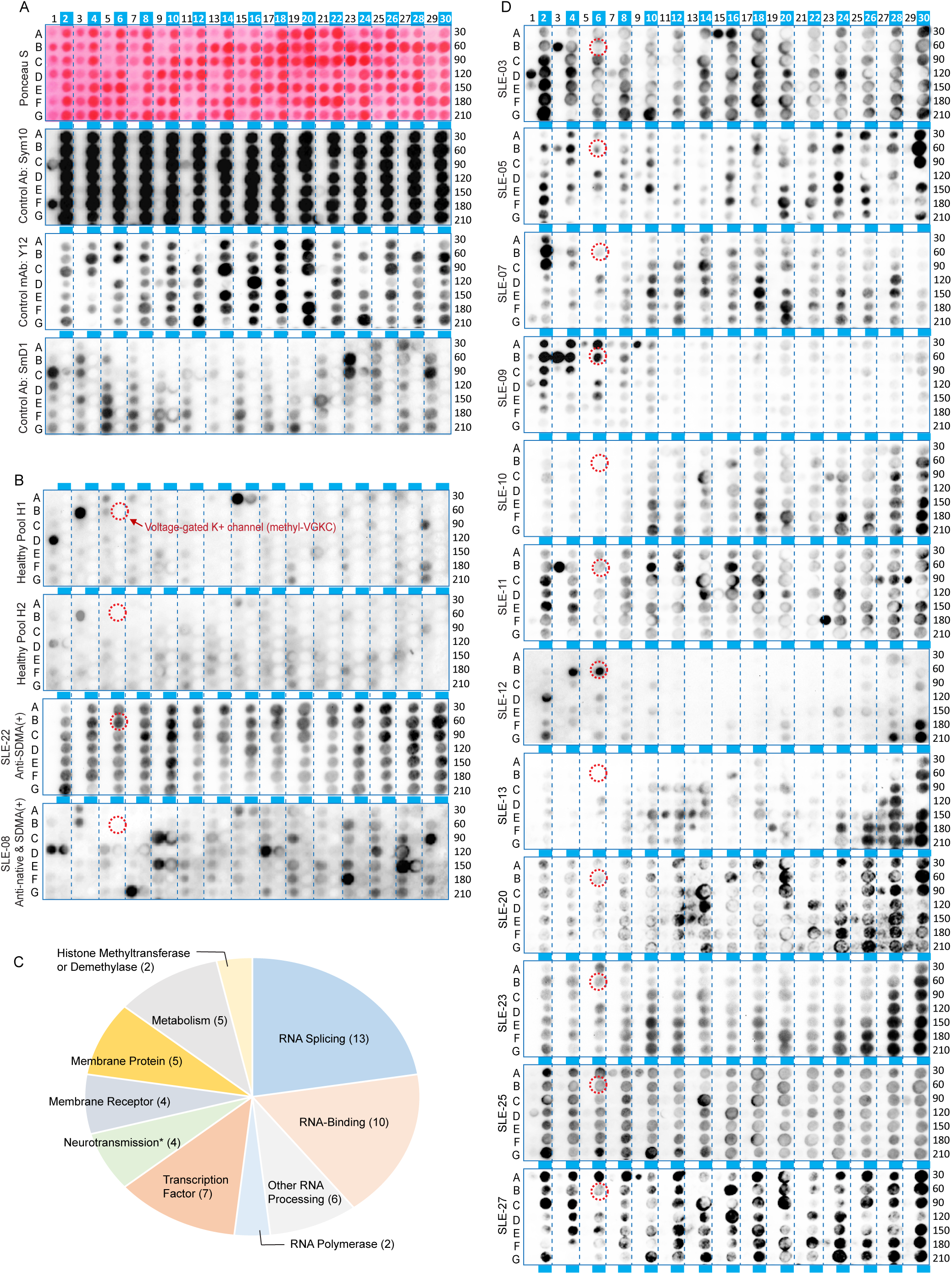
42% (13/31) of SLE patients in the discovery cohort harbor anti-SDMA autoantibodies with distinct sequence-specificity patterns. (A) A new array was generated using the same peptide sequences from the human methylarginine proteome shown in Figures 2E and 2F. The array contained adjacent peptide pairs with identical amino acid sequences: odd columns = native arginine; even columns = SDMA-containing (highlighted with blue bars). For quality control, the array was probed with Ponceau S (general peptide visualization), polyclonal SYM10 (detects SDMA peptides), monoclonal Y12 antibody (also detects SDMA peptides but with restricted selectivity), and an SmD1 antibody (recognizes a subset of native peptides, likely due to shared epitopes such as Arg-Gly repeats with SmD1). (B) The same array was probed with plasma from healthy controls H1 and H2 (also used in Figure 1A), as well as from SLE-22 and SLE-08. Healthy controls and SLE-08 reacted with a small subset of native peptides (in odd-numbered columns). In contrast, SLE-22 reacted exclusively with SDMA-containing peptides (in even-numbered columns), showing a broad pattern but with varying signal intensities across reactive peptides. The circled spot corresponds to an SDMA-containing peptide from the Voltage-Gated Potassium Channel (VGKC) protein, which showed antibody signals with SLE-22. The same methyl-VGKC peptide is also highlighted in panel D to facilitate comparison of differential patient reactivity. (C) Compilation of the methylarginine proteome (Summarized in Supplemental Table S1) and functional categorization of the corresponding proteins show that the majority are involved in RNA biogenesis, as expected. However, additional proteins include antibody accessible membrane proteins and proteins involved in neurotransmission (e.g., VGKC), suggesting they might be targeted by anti-SDMA autoantibodies on intact cells. (D) Anti-SDMA specificities among the remaining 12 SLE patients were compared using the same peptide array. Distinct antibody reactivity against the methyl-VGKC peptide (circled) was observed only in SLE-09 and SLE-12 (in addition to SLE-22 in panel B). More broadly, the 12 patients exhibited marked differences in their overwhelmingly SDMA-specific signals across the peptide panel, with only a few native peptides showing reactivity.

In addition, we tested two plasma samples from healthy donors. One showed reactivity to several native peptides, while the other exhibited no detectable antibody signals (Figure 3B). This pattern stood in stark contrast to that of SLE-22, which reacted to the majority of SDMA-modified peptides on the array, albeit with varying intensities. Intriguingly, SLE-08 reacted to approximately a dozen native peptides, most of which were distinct from those recognized by the reactive healthy control. These findings suggest that the SDMA-specific reactivity observed in SLE-22 represents only a subset of autoantibody specificities present in SLE.

Although the peptide panel representing the human methylome is enriched with proteins involved in RNA biogenesis (Figure 3C and Supplemental Table S1), it also contains unexpected membrane proteins, including those with functions in neurotransmission. Given the prevalence of neuropsychiatric lupus (NPSLE) among patients(37, 38), we investigated whether the peptide reactivity patterns could indicate cross-reactivity by SDMA-directed antibodies. We next probed the array with plasma samples from all 31 SLE patients and 8 healthy controls. Thirteen of the 31 SLE patients (42%) exhibited SDMA-specific antibody reactivity (Table 1). Among these 13 patients, the vast majority (>98%) of reactive peptides were SDMA-modified, with only a few incidental spots corresponding to native peptides (Figure 3D), in contrast to the remaining 18 patients, who showed no antibody signals to SDMA peptides (Supplemental Figure S4). Strikingly, although each of these patient samples (similar to SLE-22 in Figure 3B) reacted with only a subset of the 105 SDMA peptides, their reactivity patterns were highly distinct from one another (Figure 3D). All 13 patients displayed strict selectivity for SDMA and were female (13 of 28 female patients), whereas none of the male patients (0 of 3) tested positive (Table 1).

Consider the aforementioned membrane proteins as an example: the peptide at spot 6B on the array corresponds to an SDMA-modified peptide from the voltage-gated potassium channel (VGKC) protein, which is known to be a target of autoantibodies in autoimmune encephalitis(39, 40). Our SLE patients showed differential responses to this peptide, with the strongest reactivity observed in SLE-09, SLE-12, and SLE-22 (Figure 3B and 3D). Overall, we observed distinct antigen-reactive patterns of anti-SDMA specificity among SLE patients.

### Lupus-Prone MRL-lpr Mice Spontaneously Develop Anti-SDMA Autoantibodies

Given that variability in anti-SDMA specificity in our cohort might be influenced by genetic or ancestry factors, we turned to a syngeneic lupus mouse model, the MRL/MpJ-Faslpr/J mouse (MRL-lpr)(41). Longitudinal serum samples (n=12) were collected between 8 and 13 weeks of age, during which all mice developed anti-SDMA autoantibodies, as demonstrated by ELISA using a pair of native versus SDMA-modified 10xRG-repeat peptides (Figure 4A) or by immunoblotting using total cell lysates harvested from HEp-2 cells pretreated or not with PRMT5 inhibitor (Figure 4B). Notably, although ELISA signals against the SDMA peptide increased across the majority MRL-lpr mice, immunoblotting revealed distinct band patterns among individual syngeneic mice (Figure 4C), suggesting that anti-SDMA recognition of its carrier proteins occurs stochastically.

**Figure 4.**
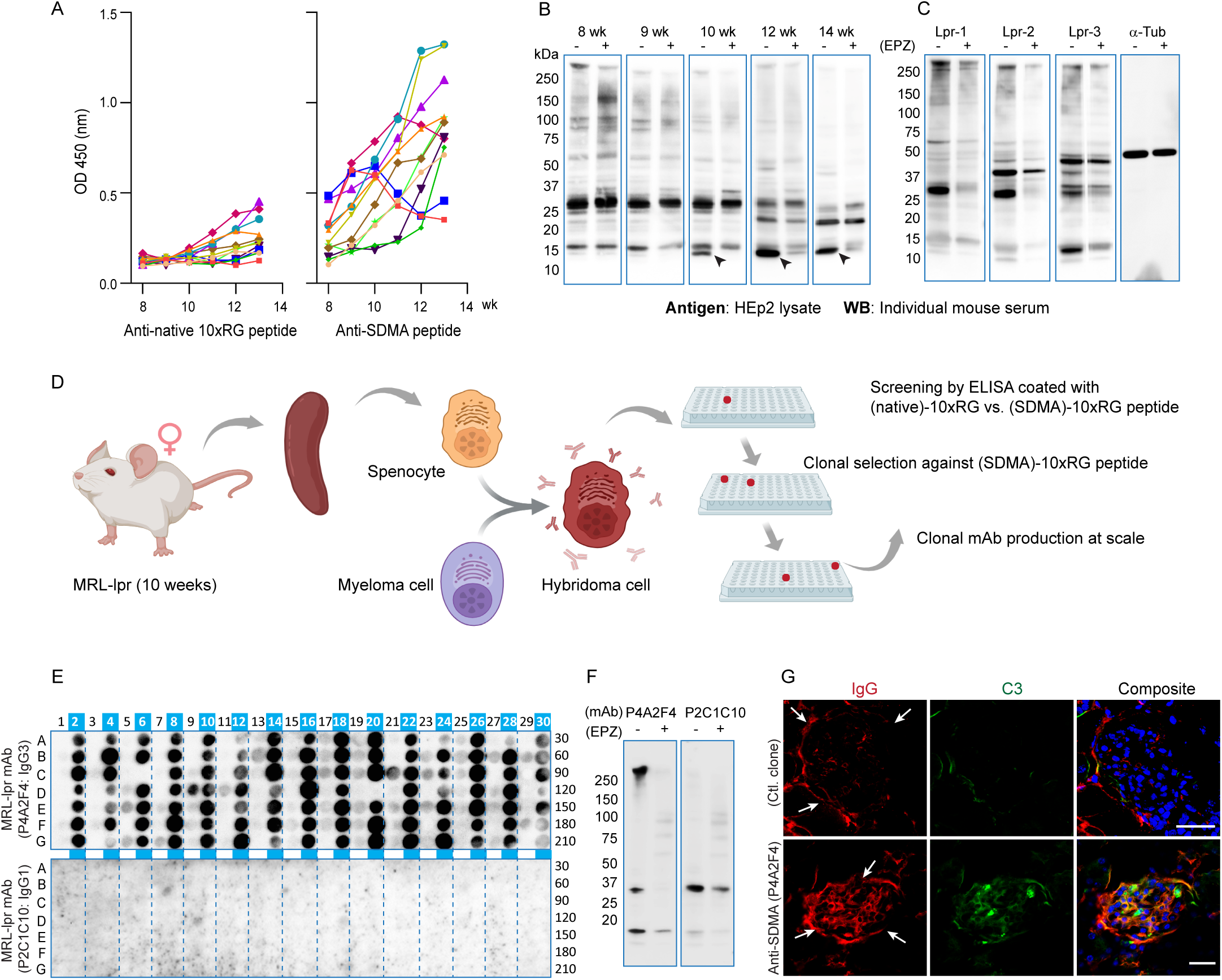
Spontaneous development of anti SDMA autoantibodies in lupus prone MRL lpr mice. (A) Sera from MRL lpr mice (n=12, bled weekly from 8 to 13 weeks of age) were analyzed by ELISA using synthetic 10×RG peptides: native sequence (GRGRGRGRGRGRGRGRGRGR) vs. SDMA-modified (lowercase ‘r’ denotes SDMA; GrGrGrGRGRGrGrGrGRGR). All 12 mice developed anti SDMA antibody titers above the corresponding native controls, and the majority (9/12) showed a gradual increase in anti-SDMA titers over time. (B) Immunoblotting of HEp-2 lysates revealed progressive changes in band patterns from 10 weeks onward. To identify SDMA specific reactive proteins, cells were pretreated with the methyltransferase inhibitor EPZ015666 (EPZ) before lysis. A prominent ∼15 kDa band (arrowhead) was markedly reduced by EPZ pretreatment, supporting its recognition by anti SDMA antibodies. (C) Sera from three 12 week old MRL lpr littermates (Lpr 1, 2, 3) each showed distinct band patterns against the cell lysates. EPZ pretreatment of the cells reduced the intensity of several bands in all three mice. (D) Workflow for isolating hybridoma clones from individual plasma cells to obtain monoclonal antibodies (mAbs) specific for SDMA. MRL lpr mice were not immunized with any antigen; thus, the resulting mAbs represent naturally arising autoantibodies. (E) Two antibody clones were tested against the native versus SDMA peptide array shown in Figurè3. Clone P4A2F4 reacted extensively with most SDMA containing peptides, whereas clone P2C1C10 showed no reactivity. (F) The two mAbs produced different band patterns on cell lysates. P4A2F4 recognized bands that were reduced in EPZ pretreated lysates, consistent with SDMA specificity. (G) After implantation of the SDMA-reactive hybridoma clone into the peritoneal cavity of healthy BALB/c mice, the kidneys showed extensive IgG deposition, reminiscent of human lupus nephritis.

Leveraging the high prevalence of anti-SDMA autoantibodies in MRL-lpr mice, we sought to further explore their clonal basis to address two key questions: whether a single anti-SDMA clone can target a broad range of autoantigens, and whether clonal specificity for SDMA-flanking sequences contributes to distinct tissue or organ involvement in individuals with SLE. To this end, we subjected naïve female ten-week-old MRL-lpr mice to a hybridoma cloning procedure (Figure 4D). Following several rounds of ELISA screening against the pair of native versus SDMA-modified peptides used in Figure 4A, we successfully isolated several SDMA-specific hybridoma clones. A representative example is shown in Figure 4E, featuring clone P4A2F4, which was used to probe the peptide array and cell lysates, in comparison with the negative control clone P2C1C10, which exhibited no reactivity against the peptide set. The array confirmed selectivity for SDMA. Notably, this monoclonal antibody exhibited broad reactivity to most SDMA-modified peptides on the array, suggesting that a single anti-SDMA autoantibody clone can potentially react to a wide range of autoantigens. Additional hybridoma clones from the mice showed both shared and distinct SDMA-specific peptide reactivity patterns on the array to varying degrees (Supplemental Figure S5), reinforcing the notion that clonal anti-SDMA antibodies can target multiple antigens. Immunoblotting against cell lysate harvested from PRMT inhibitor-treated cells further confirmed that monoclonal antibody P4A2F4 targets arginine-methylated proteins (Figure 4F). Immunofluorescence staining of cells with this antibody revealed ANA reactivity at the nuclear periphery (Figure 1C), in contrast to other isolated monoclonal antibodies (Figure 1C). Collectively, these results from the patient cohort and MRL-lpr mice suggest clonal anti-SDMA autoantibodies may be responsible for targeting a wide range of autoantigens via their shared methylarginine-containing epitopes.

### SLE Autoantibodies Cross-React with the SDMA-Modified EBNA-1 Viral Antigen of Epstein-Barr Virus

Given the repetitive nature of RG sequences and their broad presence in cellular proteins, we further explored their mechanistic link to SLE pathogenesis. Epstein-Barr virus (EBV) is widely considered a potential trigger for SLE(42, 43), with various mechanisms proposed, particularly involving its EBNA proteins(44). We examined two prominent RG-repeat sequences in EBNA1 (Figure 2A). The longer of the two spans amino acids 378-426 and contains 14 RG repeats, for which we generated a set of overlapping peptides (Figure 2H). As before, these peptides were synthesized with arginine residues either in their native form or modified as MMA, SDMA, or ADMA on a membrane array. This EBNA1-derived array was then probed with plasma from SLE patient 22 (SLE-22). The results revealed strong reactivity to all SDMA-modified peptides (Figure 2H), whereas no signals were detected for the corresponding native, MMA, or ADMA peptides. We observed similar patterns of patient antibodies specifically targeting SDMA in 12 additional patients from the cohort (13 of 31; Supplemental Figure S6). Furthermore, we compared the full-length EBNA1 protein produced in non-methylating E. coli with that produced in human HEK293 cells, which support natural arginine methylation. Immunoblotting against cell lysate with SLE plasma samples and a monoclonal antibody isolated from the lupus mouse model showed autoantibodies specifically targeting methylated EBNA1 (Figure 2I). These findings support the hypothesis that lupus autoantibodies targeting SDMA epitopes may cross-react with similar RG-repeat sequences, suggesting a new mechanism by which EBV may contribute to SLE onset.

### Anti-SDMA Autoantibodies Promote the Assembly of High-order Immune Complexes by Binding to Multiply Methylated RG Motifs on Antigens

Another aspect of the SDMA-centric hypothesis is that for multiply methylated sequences, antigen epitopes may be presented in proximity to one another and simultaneously bind multiple antibodies, thereby eliciting stronger immune responses(45). To investigate the spatial requirements for two antibody molecules to bind a pair of neighboring SDMA residues, we generated a new set of 10×RG-repeat peptides containing two SDMAs separated by 1 to 11 amino acids. When probed with SLE plasma, these SDMA-containing peptides showed antibody binding. Interestingly, peptides with two SDMAs separated by one or three amino acids exhibited antibody signals comparable to those of peptides with only a single SDMA (Supplemental Figure S7A), suggesting that closely spaced SDMAs may cause steric hindrance for binding separate antibodies. As the distance between the two SDMAs increased, antibody signals became stronger, as observed with SLE-03, SLE-22, and SLE-25, likely reflecting simultaneous binding of two IgG molecules to the dual-SDMA peptides. These findings raise the possibility that bivalent engagement of IgG with multiply methylated RG-repeat regions on a protein may drive the assembly of large immune complexes (Supplemental Figure S7B), thereby potentially augmenting pathogenicity. In keeping with this, hybridoma-bearing mice that produced an anti-SDMA monoclonal antibody showed significant IgG deposits in the kidney glomerulus (Figure 4G), reminiscent of SLE kidney pathology.

### A Validation Cohort Reveals Anti-SDMA Autoantibodies in Association with SLE Activity

To validate our findings, we recruited a new cohort of 201 Chinese SLE patients (Table 2 and Supplemental Table S2) and quantified anti-SDMA antibody titers by ELISA, using the same pair of 10×GR-repeat peptides with or without SDMA modifications. Compared to healthy controls, the SLE cohort exhibited significantly elevated anti-SDMA antibody levels (Figure 5A), with 29% of patients showing titers above the control range. Seropositivity was observed in approximately 30% of female patients (57 of 188) versus 15% of male patients (2 of 13)(Table 2), which did not reach statistical significance, possibly due to the small male sample size. As expected, reactivities to the unmodified 10×GR antigen did not differ significantly between the two groups (Figure 5B). Overall, a higher rate of seropositivity was observed in patients admitted to the hospital for the first time on SLE diagnosis, and patients with shorter disease duration are associated with higher anti-SDMA titer and prevalence of seropositivity (Figure 5C).

**Figure 5.**
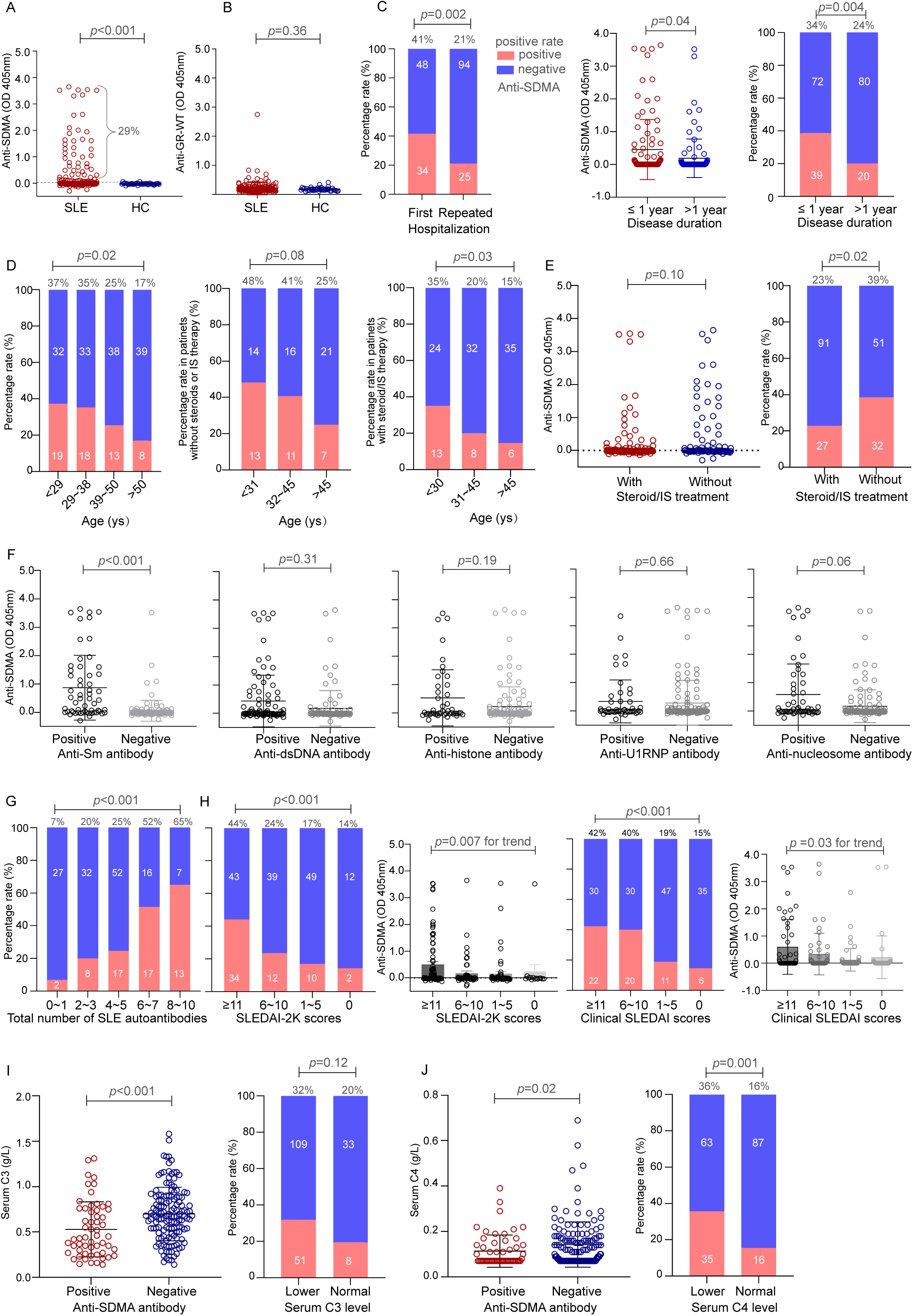
Anti SDMA titers correlate with clinical disease severity in a validation cohort of 201 SLE patients. (A,B) Plasma antibody titers against the 10xRG peptide pair either containing SDMA (A) or native (non methylated) arginine (B) were measured in 201 SLE patients and 38 healthy controls (HC). Only anti SDMA titers were significantly elevated in SLE patients vs. HC (p<0.001)(A), whereas titers against the native peptide were low and did not differ between groups (p=0.36)(B). Using the highest HC value as a cutoff (dotted line in A), the seropositivity rate for anti SDMA in SLE patients was 29%. (C) Seropositivity was higher among patients hospitalized at first diagnosis than among those with repeated hospitalizations (41% (34/82) vs. 21% (25/119), p=0.002)(left panel). Elevated anti SDMA titers correlated with a more recent SLE diagnosis, defined as less than one year since disease onset (middle panel), and patients with a disease duration of less than one year exhibited a higher prevalence of anti SDMA seropositivity than those with longer disease duration (34% vs. 24%, p=0.004)(right panel). (D) Anti SDMA seropositivity was more prevalent in younger patients: 37% (19/51) in the <29 years group vs. 17% (8/47) in the >50 years group (p for trend test=0.02)(left panel). Among patients without prior steroid or immunosuppressant (IS) treatment, 48% (13/27) of those <31 years were anti SDMA positive, compared to 25% (7/21) of those >45 years; however, the trend did not reach statistical significance (p=0.08)(middle panel). In previously treated patients, seropositivity was 35% (13/37) in the <30 years group vs. 15% (6/41) in the >45 years group (p for trend test =0.03)(right panel). (E) No significant difference in anti SDMA titers was observed between treated and untreated groups (p=0.10)(left panel), and the overall anti SDMA seropositivity was higher in untreated patients than in those receiving steroids/IS (39% (32/83) vs. 23% (27/118), p=0.02)(right panel). (F) Anti SDMA titers were significantly higher in anti Sm positive patients (p<0.001). Meanwhile, no significant associations were found between anti SDMA and seropositivity for anti dsDNA (p=0.31), anti histone (p=0.19), anti U1 snRNP (p=0.66), or anti nucleosome (p=0.06). (G) Anti SDMA seropositivity correlated significantly with the total number of SLE autoantibodies (p<0.001 for trend test). (H) By SLEDAI 2K disease severity score, anti SDMA positivity was observed in 44% (34/77) of the ≥11 group, 24% (12/51) of the 6 10 group, 17% (10/59) of the 1 5 group, and 14% (2/14) of the 0 group (p<0.001 for trend test). Anti SDMA titers also correlated positively with SLEDAI 2K scores (p=0.007). Following the exclusion of SLEDAI 2K points derived from anti dsDNA antibody titers and serum C3/C4 levels, the remaining clinical SLEDAI scores continued to demonstrate a positive correlation between anti SDMA seropositivity and SLE disease activity (p<0.001). The clinical SLEDAI score showed a significant positive correlation with anti SDMA antibody titers (p=0.03, test for trend). (I) The anti SDMA positive group had significantly lower serum C3 levels than the negative group (p<0.001), and patients with below normal C3 showed 32% anti SDMA positivity (51/160) vs. 20% (8/41) in those with normal C3; however, the trend did not reach statistical significance (p=0.12)(left panels). (J) The anti SDMA positive group had significantly lower serum C4 levels (p=0.02). Among patients with below normal C4, 36% (35/98) were anti SDMA positive, compared to 16% (16/103) with normal C4 (p=0.001)(right panels).

**Table 2:**
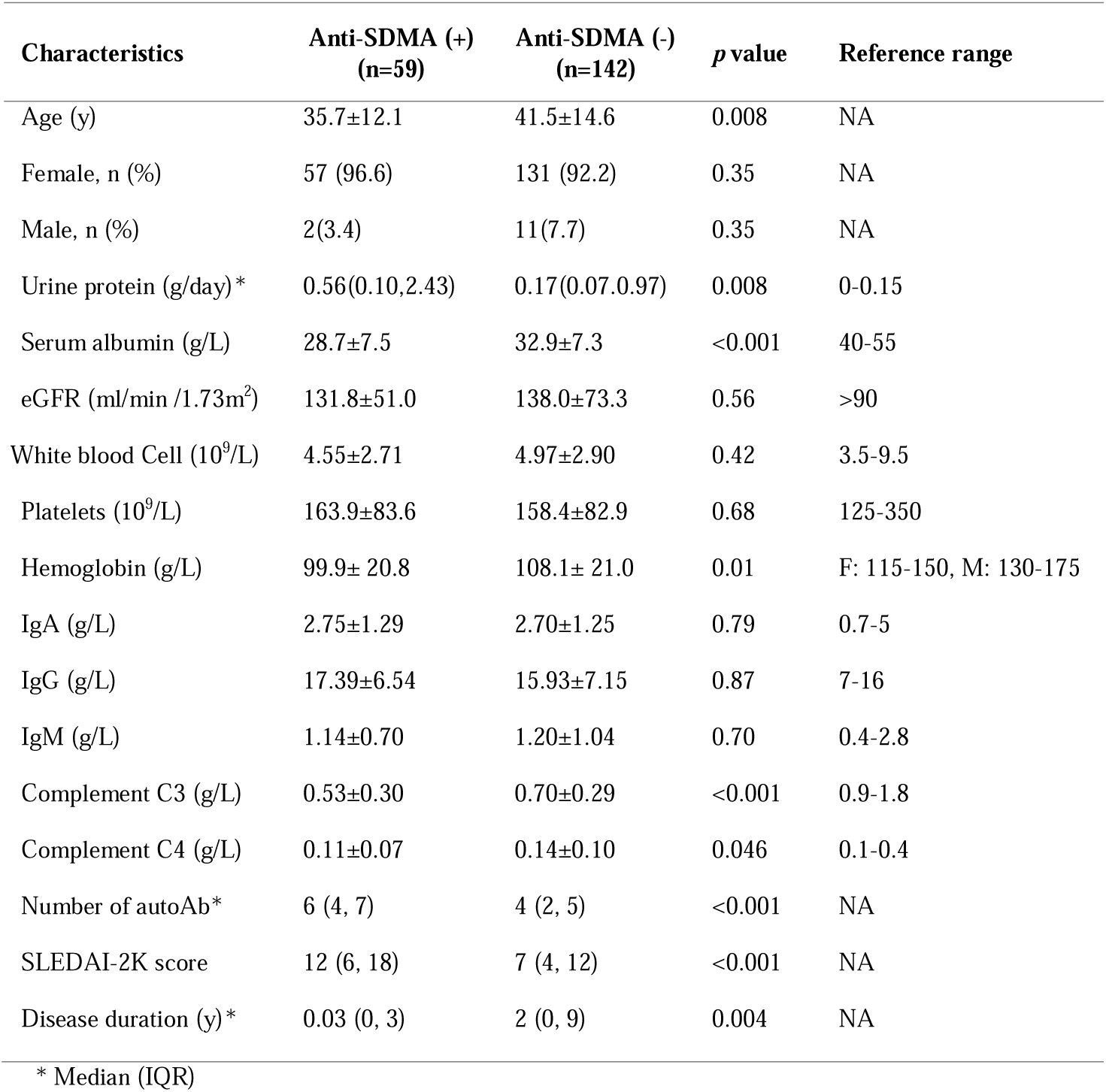
Baseline Clinical and Laboratory Features of the Validation Cohort (n=201)

| Characteristics | Anti-SDMA (+)<br>(n=59) | Anti-SDMA (-)<br>(n=142) | *p* value | Reference range |
| --- | --- | --- | --- | --- |
| Age (y) | 35.7±12.1 | 41.5±14.6 | 0.008 | NA |
| Female, n (%) | 57 (96.6) | 131 (92.2) | 0.35 | NA |
| Male, n (%) | 2(3.4) | 11(7.7) | 0.35 | NA |
| Urine protein (g/day)* | 0.56(0.10,2.43) | 0.17(0.07,0.97) | 0.008 | 0-0.15 |
| Serum albumin (g/L) | 28.7±7.5 | 32.9±7.3 | <0.001 | 40-55 |
| eGFR (ml/min /1.73m<sup>2</sup>) | 131.8±51.0 | 138.0±73.3 | 0.56 | >90 |
| White blood Cell (10<sup>9</sup>/L) | 4.55±2.71 | 4.97±2.90 | 0.42 | 3.5-9.5 |
| Platelets (10<sup>9</sup>/L) | 163.9±83.6 | 158.4±82.9 | 0.68 | 125-350 |
| Hemoglobin (g/L) | 99.9± 20.8 | 108.1± 21.0 | 0.01 | F: 115-150, M: 130-175 |
| IgA (g/L) | 2.75±1.29 | 2.70±1.25 | 0.79 | 0.7-5 |
| IgG (g/L) | 17.39±6.54 | 15.93±7.15 | 0.87 | 7-16 |
| IgM (g/L) | 1.14±0.70 | 1.20±1.04 | 0.70 | 0.4-2.8 |
| Complement C3 (g/L) | 0.53±0.30 | 0.70±0.29 | <0.001 | 0.9-1.8 |
| Complement C4 (g/L) | 0.11±0.07 | 0.14±0.10 | 0.046 | 0.1-0.4 |
| Number of autoAb* | 6 (4, 7) | 4 (2, 5) | <0.001 | NA |
| SLEDAI-2K score | 12 (6, 18) | 7 (4, 12) | <0.001 | NA |
| Disease duration (y)* | 0.03 (0, 3) | 2 (0, 9) | 0.004 | NA |

\* Median (IQR)

Having validated that this invariant sequence with SDMA modifications broadly captures anti-SDMA activities in a subset of SLE patients, we stratified approximately equal numbers of patients into four age groups: <29, 29-38, 39-50, and >50 years. A clear trend of anti-SDMA positivity emerged in younger patients, with rates of 37%, 35%, 25%, and 17%, respectively (Figure 5D). To control for the potential confounding effect of treatment history on antibody positivity, we analyzed patients with (n=118) or without (n=83) prior exposure to steroids or immunosuppressants, dividing them equally into three age groups. The inverse correlation between anti-SDMA positivity and age persisted; the youngest untreated group showed 48% antibody positivity, followed by 41% in the middle age group and 25% in the oldest group (Figure 5D). Consistent results were observed in the subgroup of patients undergoing immunosuppression (Figure 5E). Notably, no significant differences in anti-SDMA titers were found between treated and untreated individuals when analyzed as two separate groups, although the anti-SDMA seropositivity rate was higher in untreated patients.

We next examined whether anti-SDMA antibodies correlate with conventional SLE autoantibody tests. Patients were stratified into autoantibody-positive and -negative groups for several specificities, including anti-Sm, anti-dsDNA, anti-histone, anti-U1 snRNP, and anti-nucleosome (Figure 5F). Anti-SDMA levels were significantly higher in anti-Sm-positive patients than in anti-Sm-negative patients (0.87 vs. 0.06, p<0.001), consistent with the presence of SDMA-modified RG dipeptide sequences in Sm antigens (Supplemental Figure S8). While anti-SDMA levels are generally higher among patients who tested positive for other SLE autoantibodies, the trend did not reach statistical significance (Figure 5F), likely reflecting the involvement of different antigenic epitopes. Notably, patients with a greater number of positive autoantibodies tended to have a higher rate of anti-SDMA seropositivity (Figure 5G and Supplemental Table S2), suggesting that anti-SDMA antibodies partially mirror the broader autoimmune activity in SLE.

### Anti-SDMA Autoantibody Titer Correlates with SLE Disease Activity

A key distinction between the synthetic SDMA-10×RG antigen and natural Sm antigens is that the latter are derived from human specimens, leaving their degree of SDMA modification unknown. Because our peptide antigen contains well-defined modifications on multiple arginine residues and is paired with an unmodified control, it may be better suited for detecting SDMA-specific reactivities. We next investigated whether anti-SDMA levels alone reflect SLE disease activity. Patients were stratified into five groups based on SLEDAI-2K scores: 0, 1-5, 6-10, and ≥11. Higher SLEDAI-2K score groups generally showed a greater proportion of anti-SDMA-positive patients: 14%, 17%, 24%, and 44%, respectively (Figure 5H). Among anti-SDMA-positive cases, antibody titers also correlated with SLEDAI-2K scores, with the highest disease activity group displaying particularly elevated titers. Furthermore, to eliminate the contributions of serology measurements to SLEDAI-2K, we recalculated clinical SLEDAI scores after removing points attributable to anti-dsDNA and complement C3 and C4 levels. A similar correlation between anti-SDMA and clinical SLEDAI was observed, both in terms of antibody prevalence and titer (Figure 5H). Additionally, the anti-SDMA-positive group had significantly lower serum C3 and C4 levels (Figure 5I-5J, and Table 2), indicating greater clinical severity than the antibody-negative group. Regarding organ involvement, seropositivity and titer showed a trend toward correlation with kidney and hematologic involvement (Supplemental Figure S9A-S9J). In contrast, SLE-related pulmonary, cardiac, neuropsychiatric, and joint manifestations were not significantly correlated with anti-SDMA seropositivity (Supplemental Figure S9K-S9N).

## Discussion

Our study examined the prevalence of anti-SDMA autoantibodies in SLE and their implications for pathogenic mechanisms, clinical presentation, and disease activity. We found that anti-SDMA autoantibodies were present in 30-40% of SLE patients, with a clear trend toward higher prevalence among younger patients. Plasma anti-SDMA titers correlated with disease activity as measured by SLEDAI-2K, as well as with markers of complement activation, renal involvement, and hematologic abnormalities.

Mechanistically, although anti-SDMA seropositivity partially overlaps with certain traditional SLE autoantibodies, particularly anti-Sm antigen positivity, we discovered that these autoantibodies specifically target posttranslational modifications involving symmetric dimethylation of arginine residues located within sequences rich in RG dipeptide repeats. Proteomic analysis further revealed that these RG motifs frequently reside in proteins involved in RNA biogenesis, notably ribonucleoproteins, many of which are already established SLE autoantigens. Intriguingly, our discovery study of 31 SLE patients, which utilized a panel of 105 SDMA-containing peptides representing the human protein methylome alongside their corresponding native sequences as controls, demonstrated not only that patient antibodies exhibit exclusive specificity for methylated epitopes but also that the SDMA residues and their flanking amino acid sequences jointly dictate autoantibody recognition. Notably, no two anti-SDMA-positive SLE patients shared identical patterns of peptide reactivity, suggesting that anti-SDMA autoantibodies can target a diverse array of proteins, with specificity varying across individual patients. Given the strong correlation between anti Sm and specific anti SDMA reactivities, it is plausible that clinically detected anti Sm positivity serves as a proxy for a broader anti SDMA response in SLE patients.

The observed reactivity of SLE autoantibodies across diverse SDMA-modified epitopes prompted us to investigate whether clonal SLE autoantibodies could be responsible for broad reactivity to a panel of lupus antigens through shared recognition of arginine-methylated epitopes. To address this, we isolated individual hybridoma clones from MRL-lpr mice and analyzed each clone against our panel of 105 SDMA-containing peptides. Remarkably, a single monoclonal antibody derived from an unimmunized MRL-lpr mouse reacted broadly with a wide range of SDMA peptides. Nevertheless, individual monoclonal antibodies exhibited unique profiles of both peptide reactivity and antinuclear antigen patterns. Taken together, these observations raise the possibility that clonal SLE autoantibodies against arginine methylated Sm/RNP epitopes are stochastically generated, with recognition biased toward a flanking consensus motif, a feature that could contribute to their cross reactivity with other antigenic targets.

We acknowledge several limitations in this study, particularly that our key findings on epitope specificity derive from synthetic antigen array data and hybridoma clone reactivity. A major caveat is the markedly higher antigen density on this array compared to standard assays (e.g., HEp 2 cell based or whole protein antigen tests). Whether the array signals genuinely represent immune responses against native endogenous and viral antigens therefore remains an open question that requires additional validation. Our SDMA-centric model of SLE autoantibodies further predicts potential cross-reactivity with the Epstein-Barr virus protein EBNA-1, which contains two RG-rich segments. We demonstrated that methylated EBNA-1 is recognized by SDMA-reactive plasma samples from SLE patients, suggesting a possible mechanism by which EBV may contribute to SLE pathogenesis. This finding aligns with other proposed mechanisms linking EBV to lupus development(44, 46). Notably, EBNA 2 also contains a seven RG repeat segment known to be methylated(47). Conversely, we speculate that pan-specific anti-SDMA antibodies could provide inadvertent antiviral benefits against EBV or other viruses harboring methylated RG-rich epitopes.

The RG-rich segments found in RNA-binding proteins have recently been shown to mediate RNA-induced liquid-liquid phase separation (LLPS) and stress granule association(24, 48, 49). Arginine methylation fine-tunes this liquid-phase homeostasis by suppressing RG-driven phase separation(26, 50). Given that LLPS of viral nucleocapsid proteins and other components is critical for virion assembly within infected host cells, further in-depth studies are warranted to investigate the genetic—and potentially evolutionary—basis of anti-SDMA antibodies as an innate antiviral defense mechanism that may also predispose individuals to autoimmunity. This holds true even though host autoantigens are generally concealed within cells and are released only upon cell death.

Given the varying lengths and mixed amino acid compositions of RG-rich segments across different proteins, heterogeneity in the methylation levels of these RG repeats is expected. A unique feature of antigens containing multiple SDMA residues is their ability to simultaneously bind multiple antibodies, thereby forming supersized, lattice-like immune complexes. Whether this property of anti-SDMA-directed SLE contributes to high levels of tissue deposition remains unclear. Future studies at more granular levels—examining pathological and serological correlations, as well as the spectrum of flanking sequences that constitute SDMA epitopes in individual patients—are warranted.

## Methods

### Sex as a biological variable

Both female and male participants were included in the patient and healthy control cohorts. In animal study, only female MRL-lpr mice were used because SLE predominantly affects females and disease develops earlier and more severely in female mice.

### Study cohorts and ethical approval

The initial discovery cohort consisted of 31 SLE patients recruited at Northwestern Memorial Hospital in 2015. A separate validation cohort of 201 SLE patients was enrolled at the First Affiliated Hospital of Xi’an Jiaotong University between May 2019 and December 2022. In both cohorts, SLE was diagnosed according to the EULAR/ACR criteria, based on clinical and laboratory findings after excluding alternative diagnoses. For the validation cohort, clinical data were collected, including age, sex, proteinuria, serum albumin, serum creatinine, estimated glomerular filtration rate (eGFR), IgA, IgG, IgM, complement C3 and C4, autoantibody spectrum, complete blood count, and final diagnosis. Renal involvement was defined based on the 2019 EULAR/ACR criteria, including proteinuria ≥0.5 g/24h (or equivalent urine protein-to-creatinine ratio ≥0.5 g/g) or biopsy-proven lupus nephritis (Class II-V). Plasma samples were collected and stored at −80°C until use. The studies were conducted in accordance with the Declaration of Helsinki and were approved by the Medical Ethics Committee of Northwestern University (IRB: STU00105534) and the First Affiliated Hospital of Xi’an Jiaotong University (IRB: XJTU1AF2026LSYY-0156).

### Cell culture

Human epithelial type 2 (HEp-2) cell line (ATCC, CCL-23) was used as the antigen source for immunoblot detection of autoantibodies in plasma samples. Cells were cultured in Dulbecco’s Modified Eagle Medium supplemented with 10% fetal bovine serum. To suppress cellular protein SDMA modification, cells were treated with 25 µM of PRMT5 inhibitor EPZ015666 (MilliporeSigma, SML1421) for 5 days. To prepare cell lysates, cells were cultured in 10-cm tissue culture dishes to 90-100% confluence, then washed three times with ice-cold phosphate-buffered saline (PBS) and lysed in 1 mL of 2x Laemmli sample buffer (Bio-Rad Laboratories, 1610737), followed by brief sonication to reduce viscosity. Insoluble material was removed by centrifugation at 16,000 x g for 15 min, and the supernatant was collected as total cell lysate.

### Western blot and immunofluorescence

For immunoblotting, equal volumes of HEp-2 cell lysate (15 µL per lane) or recombinant proteins (20 ng/well) were supplemented with Tris-(2-Carboxyethyl) phosphine (TCEP) to a final concentration of 40 mM, heated at 95°C for 2 min, separated by SDS-PAGE on 10% gels, and transferred onto 0.2-µm polyvinylidene difluoride (PVDF) membranes (Cytiva, 10600021). Membranes were blocked with 5% non-fat milk in Tris-buffered saline containing 0.1% Tween-20 (TBST) for 1 h at room temperature, followed by incubation overnight at 4°C with human plasma samples (1:300 dilution), mouse plasma samples (1:300 dilution), α-Tubulin antibody (SantaCruz, sc-32293; 1:500 dilution) or FLAG-M2 antibody (MilliporeSigma, F1804; 1:1000 dilution) diluted in blocking buffer. After washing with TBST, membranes were incubated for 2 h at room temperature with horseradish peroxidase (HRP)-conjugated anti-human IgG (SouthernBiotech, 2087-05) or anti-mouse IgG (SouthernBiotech, 1038-05) secondary antibodies. Immunoreactive bands were visualized using enhanced chemiluminescence substrate (Promega Corporation, W1015) and imaged using the BioRad ChemiDoc XRS+ System.

For immunofluorescence staining, HEp-2 cells were seeded in 12-well tissue culture plates containing 18-mm circular glass coverslips and cultured to approximately 60-70% confluence. Cells were washed three times with ice-cold PBS and fixed with 4% paraformaldehyde for 10 min at room temperature. After washing with PBS, cells were permeabilized and blocked in PBS containing 0.1% Triton X-100 and 5% donkey serum for 1 h at room temperature. Human plasma samples (1:80 dilution) or laboratory-generated monoclonal antibodies (1:1000 dilution) were diluted in blocking buffer and incubated with the cells overnight at 4°C. SYM10 antibody (1:100 dilution; MilliporeSigma 07-412) was included as a co-stain where indicated. Following three washes with PBST, cells were incubated for 2 h at room temperature in the dark with Alexa Fluor 594-conjugated anti-human IgG (Jackson ImmunoResearch, 709-585-149) or Alexa Fluor 594-conjugated anti-mouse IgG (Jackson ImmunoResearch, 715-585-151), together with Alexa Fluor 488-conjugated anti-rabbit IgG (Jackson ImmunoResearch, 711-545-152). Nuclei were counterstained with DAPI for 5 min. Coverslips were mounted using antifade mounting medium and imaged using a Nikon A1 Confocal Microscope.

### Peptide array analyses

Custom 15-mer peptide sequences were synthesized directly onto arrays using a robotic Cellu-Spot system (Intavis AG, Köln, Germany) according to a programmed synthesis cycle. Synthesis employed Fmoc-protected amino acids, including methylarginine derivatives. Each array design was synthesized on two identical membranes to accommodate the large number of samples and minimize loss of assay performance from repeated membrane use. Following synthesis, the peptide array membranes were stained with Ponceau S to visualize and confirm the presence of peptide material at each spot. The arrays were subsequently probed with healthy control and SLE plasma samples from a discovery cohort, mouse serum samples, MRL-lpr mice-derived hybridoma monoclonal antibodies, and commercial antibody standards. Between successive probing cycles, the peptide array membranes were treated with stripping buffer (Thermo Fisher Scientific; Cat. No. 21059) and then reprobed with the next sample.

In each probing round, the membranes were washed three times in TBST buffer (50 mM Tris-HCl, pH 7.4, 150 mM NaCl, 0.1% Tween 20) for 5 minutes per wash, then blocked with 5% non-fat milk in TBST overnight at 4°C. The membranes were then probed overnight at 4°C with plasma (1: 1000 dilution), SYM10 antibody (MilliporeSigma, 07-412; 1:1000 dilution), ASYM24 antibody (MilliporeSigma, 07-414; 1: 500 dilution) or SNRPD1/Sm-D1 antibody (Abcam, ab79975; 1:250 dilution) diluted in 3% milk in TBST. Following three washes with TBST, a suitable HRP-conjugated secondary antibody diluted in 1% milk in TBST was incubated with the membrane for one hour. After a final five washes with TBST (5 minutes each), the membranes were developed using enhanced chemiluminescence (ECL) and visualized with a gel imaging system. The covalent linkage of peptides to the membrane enabled it to withstand the harsh stripping procedure without loss of performance, allowing reliable reprobing for dozens of rounds.

### Recombinant protein production

Recombinant Epstein-Barr nuclear antigen 1 (EBNA-1) protein was produced in both mammalian HEK293 cells and prokaryotic BL21(DE3) Escherichia coli cells. DNA fragments encoding EBNA-1 (with deletion of the Gly-Ala repeat region) and a C-terminal FLAG/6×His tag were synthesized by Integrated DNA Technologies and cloned into pcDNA3 vector for mammalian expression and pET30a vector for bacterial expression, respectively.

For mammalian expression, HEK293 cells were transfected with the pcDNA3-EBNA-1 expression plasmid using Lipofectamine™ 2000 Transfection Reagent (ThermoFisher, 11668027) following the manufacture’s instruction, and stable cell lines were established by G418 (GoldBio, G-418-10) antibiotic selection. Positive clones with confirmed EBNA-1 expression were expanded in 15-cm tissue culture dishes. Cells were lysed in RIPA buffer supplemented with protease inhibitor cocktail, and recombinant EBNA-1 protein was purified from clarified cell lysates using His Mag Sepharose™ Ni beads (Cytiva, 28967390) according to the manufacturer’s protocol. For bacterial expression, chemically competent BL21(DE3) cells were transformed with the pET30a-EBNA-1 plasmid and selected on LB agar plates containing kanamycin (50 μg/mL). Single colonies with confirmed recombinant protein expression were inoculated into LB broth containing kanamycin and cultured at 37°C until mid-log phase, followed by induction with 0.4 mM IPTG overnight at 18°C. Bacterial pellets were harvested by centrifugation and lysed by sonication. Recombinant EBNA-1 protein was purified from clarified lysates using a HisTrap HP column (Cytiva, 17524701) following the manufacturer’s instructions. The purified proteins were aliquoted and stored at −80°C until use.

### Enzyme-Linked Immunosorbent Assay (ELISA)

To measure antibody titers against SDMA-specific epitopes by ELISA, two N-terminal biotinylated peptides were synthesized by the Northwestern University Peptide Synthesis Core and used for ELISA development: Biotin-GRGRGRGRGRGRGRGRGRGR and Biotin-GrGrGrGRGRGrGrGrGRGR, in which “r” denotes SDMA. High-binding ELISA plates (Greiner Bio-One, 655081) were coated with 250 ng/well of streptavidin (Leinco Technologies, S203) overnight at 4°C. After washing five times with TBST, the two peptides diluted in TBST were added to adjacent rows in parallel at a concentration of 1 µg/mL and incubated for 1 h at room temperature. Plates were then washed five times with TBST and blocked with 5% bovine serum albumin (BSA) in TBST. For plasma samples, plasmas were diluted 1:200 in blocking buffer. For hybridoma culture supernatants, the medium was diluted 1:1 with 2x blocking buffer. The diluted samples were then added to the blocked wells and incubated overnight at 4°C. Plates were subsequently washed five times with TBST and incubated for 2 h at room temperature with horseradish peroxidase-conjugated anti-human IgG (SouthernBiotech, 2087-05) or anti-mouse IgG (SouthernBiotech, 1038-05) secondary antibody. After five additional washes with TBST, signals were developed using a TMB chromogenic ELISA substrate, and the reactions were terminated by adding 2 N HCl. Absorbance at 450 nm was measured using a microplate reader.

Anti-Epstein-Barr virus viral capsid antigen (EBV-VCA) IgG and anti-Smith (Sm) antigen IgG levels in plasma samples from the SLE discovery cohort were measured using commercial ELISA kits (ab108730, Abcam; and KA0948, Abnova, respectively). Samples were classified as positive or negative according to the manufacturers’ instructions.

### Generation of hybridomas from MRL-lpr mice

All animal experiments in this study were approved by the Northwestern University Institutional Animal Care and Use Committee (IACUC) under protocol IS00000429. Monoclonal autoantibodies were generated from lupus-prone MRL-lpr mice using conventional hybridoma technology. To identify mice with high plasma autoantibody levels against SDMA-modified antigens, plasma samples collected from individual MRL-lpr mice were first screened using the peptide ELISA described above. Samples exhibiting high reactivity toward SDMA-modified peptides were further evaluated by immunoblotting using lysates from untreated HEp-2 cells or HEp-2 cells treated with EPZ015666, which suppresses SDMA modification of nuclear antigens. Mice whose plasma samples showed stronger reactivity toward SDMA-peptide and HEp-2 cells proteins present in untreated but reduced in EPZ015666-treated HEp-2 lysates were selected as spleen donors for hybridoma generation.

Murine myeloma cell line SP2/0-Ag14 were thawed and expanded for approximately 1 week before cell fusion in RPMI-1640 medium supplemented with 20% fetal bovine serum and sodium pyruvate. One to two days before fusion, myeloma cells in logarithmic growth phase were further expanded in 175-cm² culture flasks. For feeder cell preparation, peritoneal macrophages were isolated from naïve female BALB/c mice 1 day before fusion and seeded into 96-well plates containing HAT selection medium.

Selected MRL-lpr mice were euthanized, and spleens were harvested under sterile conditions. Splenocytes were prepared by mechanical dissociation, 70-µm cell strainer filtration, and then treated with red blood cell lysis buffer. Splenocytes were then mixed with SP2/0-Ag14 cells at a 3:1 ratio. For cell fusion, mixed cells were washed with serum-free RPMI-1640 and fused with pre-warmed 50% polyethylene glycol by slow dropwise addition with gentle mixing. After gradual dilution with serum-free medium, cells were centrifuged and resuspended in HAT selection medium at approximately 2.5 × 10⁶ cells/mL, then plated onto feeder cell-containing 96-well plates at 100 µL per well.

Hybridoma cells were cultured at 37°C with 5% CO₂, and half of the HAT medium was replaced every 2-3 days until colonies became visible. HAT medium was changed to HT medium on days 7-10, followed by complete medium on day 14. Hybridoma supernatants were screened by ELISA for reactivity against SDMA-modified peptides. Positive clones were subjected to two additional rounds of subcloning to establish stable monoclonal cell lines.

The isotype of each monoclonal antibody in the hybridoma culture supernatant was determined using a mouse monoclonal antibody isotyping kit (Sigma-Aldrich, ISO2) following the manufacturer’s antigen-mediated ELISA protocol.

### Production of monoclonal autoantibodies

Selected hybridoma clones were expanded in culture and used for monoclonal antibody production in pristane-primed female BALB/c mice. Briefly, mice were intraperitoneally injected with pristane (0.4 mL/mouse), followed by intraperitoneal injection of hybridoma cells (3 x10 cells/mouse) 7 days later. Ascites fluid was collected when abdominal distension became apparent and clarified by centrifugation to remove cellular debris. Monoclonal antibodies were purified from ascites fluid using HiTrap Protein G HP column (Cytiva, 29048581) according to the manufacturer’s protocols. Purified antibodies were concentrated and buffer-exchanged into PBS using Amicon Ultra centrifugal filter units (MilliporeSigma, UFC9030), aliquoted, and stored at −80°C until use.

### Arg-Gly motif frequency search

A non-redundant set of RefSeq human proteins was screened using a sliding window of 10 consecutive amino acids, moving from the N-terminus to the C-terminus of each protein. A hit was defined as the presence of at least two Arg-Gly (RG) motifs within a single 10-amino-acid window. A weighted score based on the total number of hits was calculated for each protein. The top 382 ranked proteins by this score were subjected to gene ontology enrichment analysis (geneontology.org) for cellular component and biological process categories.

### Statistical analysis

Normally distributed continuous variables are presented as mean ± standard deviation (SD), while non-normally distributed continuous variables are presented as median with interquartile range (IQR). Between-group differences in continuous variables were assessed using the independent-samples t-test for normally distributed data and the Mann-Whitney U test for non-normally distributed data. The Jonckheere-Terpstra trend test was used to evaluate the association between SLEDAI-2K scores and anti-SDMA titers, based on the distribution and variance homogeneity of the titers. The Cochran-Armitage trend test (reported as the linear-by-linear association chi-square test) was applied to assess linear trends between ordinal independent variables, such as number of autoantibodies, age groups, or increasing SLEDAI-2K scores, and anti-SDMA seropositivity. Fisher’s exact test was used for comparisons of categorical variables between groups, such as male versus female, when the expected frequency in any cell of the 2×2 table was ≤5. A two-sided p-value < 0.05 was considered statistically significant. All analyses were performed using SPSS version 26.0.

## Data availability

Values for all data points in figures are reported in the Supporting Data Values file. Supplemental Information Supplemental Table S1-S2 Supplemental Figures S1-S9 Acknowledgements We would like to thank Dr. Cybele Ghossein and Ms. Laura J. Nishi for collecting blood samples and Dr. Shawn Li for assistance with the peptide array. We would also like to thank Drs. Tomokazu Souma, Richard M. Pope, and Rosalind Ramsey-Goldman for their advice on the study. This work was supported through grants from the National Institutes of Health (R21AI131087 and R01EB033377 to J.J.) and National Natural Science Foundation of China (Grant No. 82270747 to X.X.).

## Author Contributions

Conceptualization: P.L., X.X., J.J.

Methodology: P.L., X.X.

Investigation: P.L., H.L., A.Z.W., J.G.P., X.X., J.J.

Formal Analysis: P.L., X.X., J.J.

Funding acquisition: X.X., J.J.

Manuscript preparation: All authors contributed.

## AI Disclosure

The authors declare that no AI tools were used in the original research for or writing of this article.

## Data Statement

All data that support the findings are included within the manuscript. Other source data related to this study are available from the corresponding authors upon reasonable request.

