## Supplemental Figures for "Symmetric Dimethylarginine Modification Drives Shared and Divergent Antigen Reactivity in Lupus"

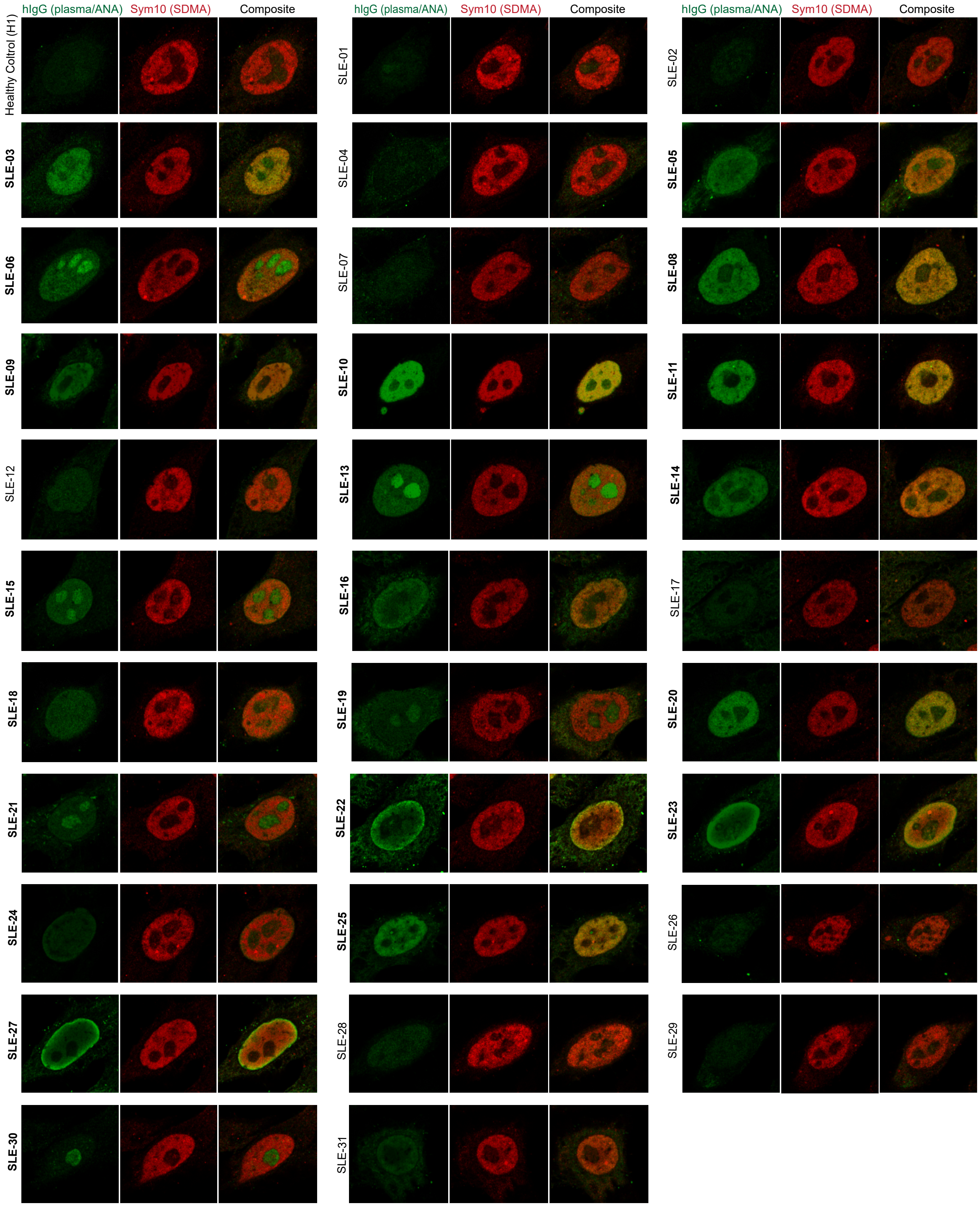


**Supplemental Figure S1. Anti-nuclear antibody (ANA) screening of 31 SLE plasma samples.**

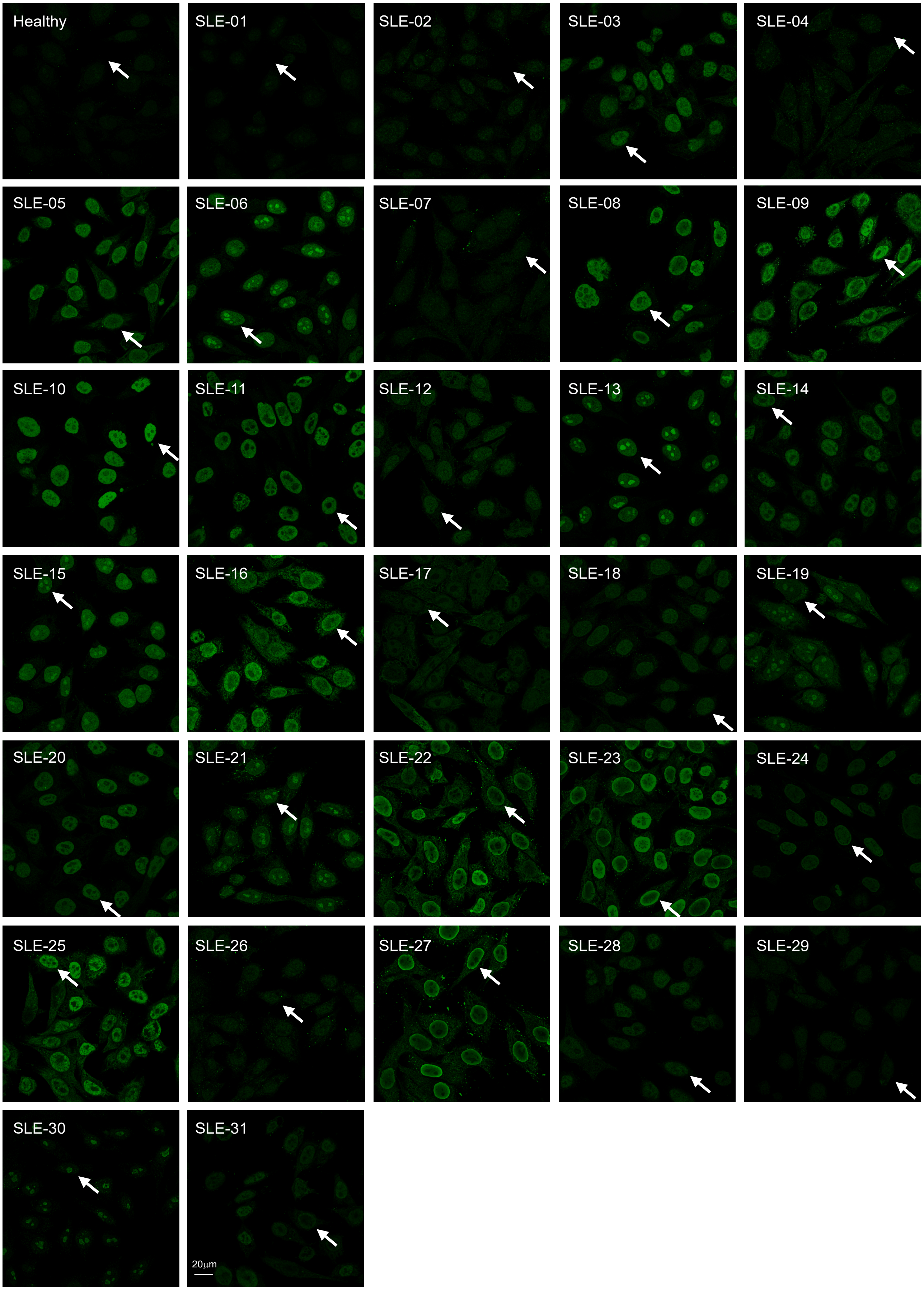


**Supplemental Figure S2. Low-magnification ANA immunofluorescence staining of HEp-2 cells with SLE patient plasma.** Plasma samples from 31 SLE patients and one healthy control were used to probe HEp-2 cells, with low-magnification images captured for 20–30 cells per sample. Arrows indicate representative single cells from each sample that are shown at higher resolution in Figure 1B and Supplemental Figure S1.

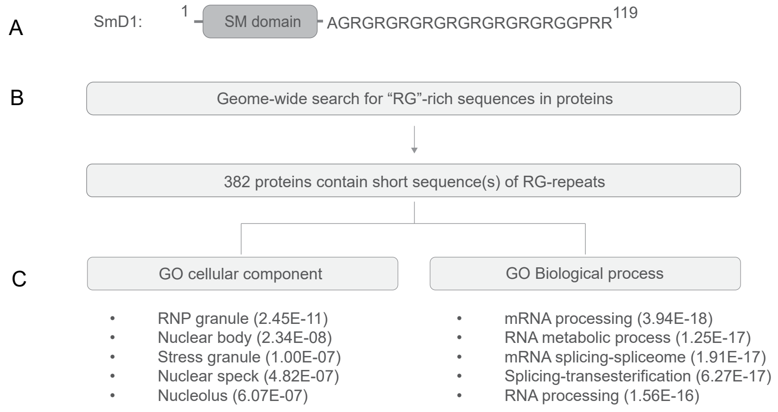


**Supplemental Figure S3. Enrichment of Arg-Gly (RG)-repeat motifs in nuclear proteins involved in RNA biogenesis.**
A. Human SmD1 protein, as an example, consists of an N-terminal SM domain and a C-terminus segment populated with RG repeats.
B. Genome-wide search for proteins containing at least two RG motifs within eight amino acids of each other yielded a total of 382 proteins.
C. Gene ontology analyses revealed their enrichment among nuclear proteins with functions in RNA biogenesis. The p-value reflects the probability that the observed overlap between a given gene list and a specific biological category is due to chance.

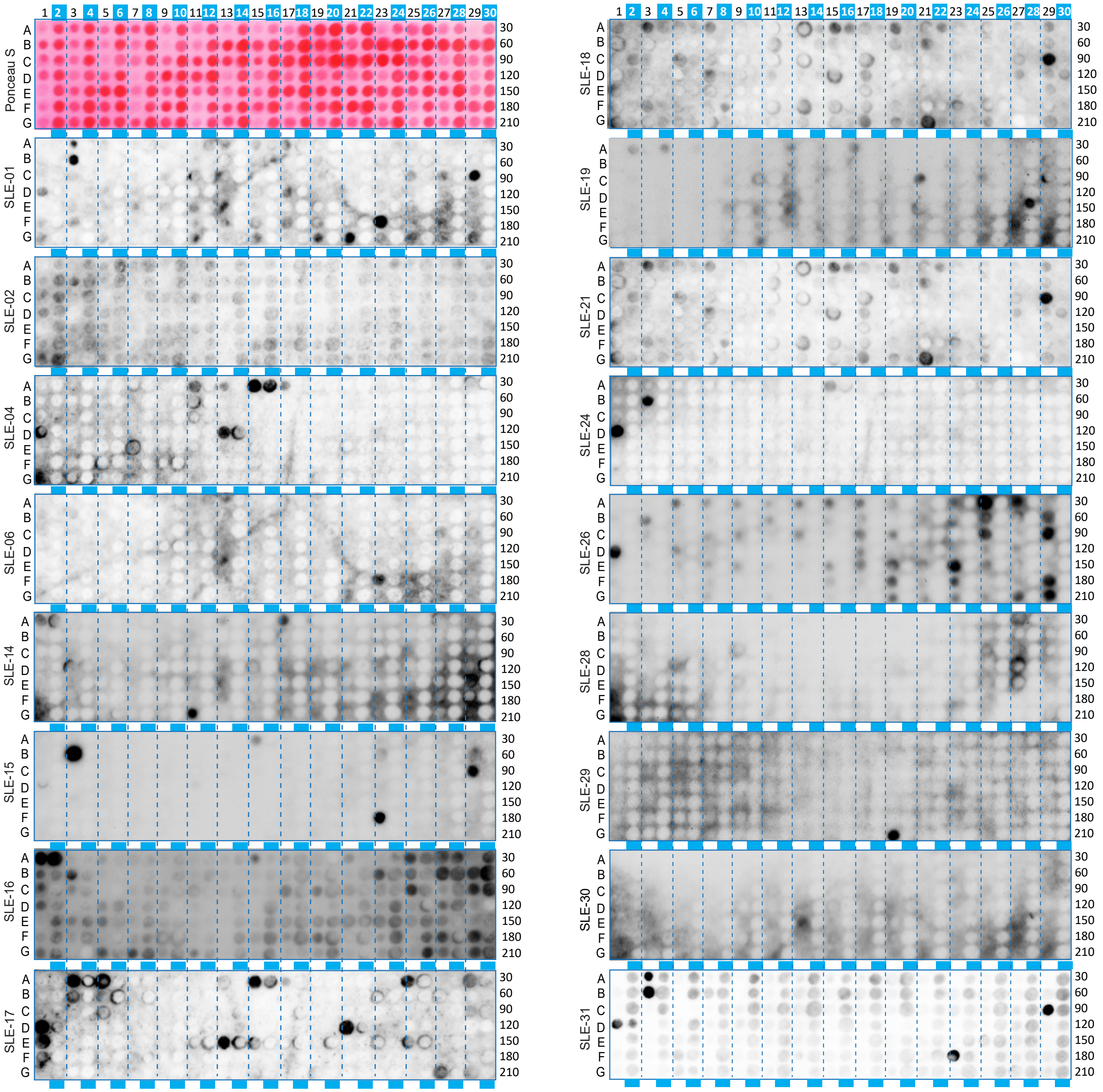

**Supplemental Figure S4. Array results of eighteen anti-SDMA-negative patients in the discovery cohort.**
The top-left panel displays the overall array layout visualized by Ponceau S staining. Odd columns contain unmodified peptides; even columns (blue bars) contain peptides in which all arginine residues have been substituted with symmetric dimethylarginine (SDMA). Arrays were probed with plasma from all 31 SLE patients in the discovery cohort. Shown here are samples with either no antibody-reactive peptides or antibodies binding predominantly to unmodified peptides.

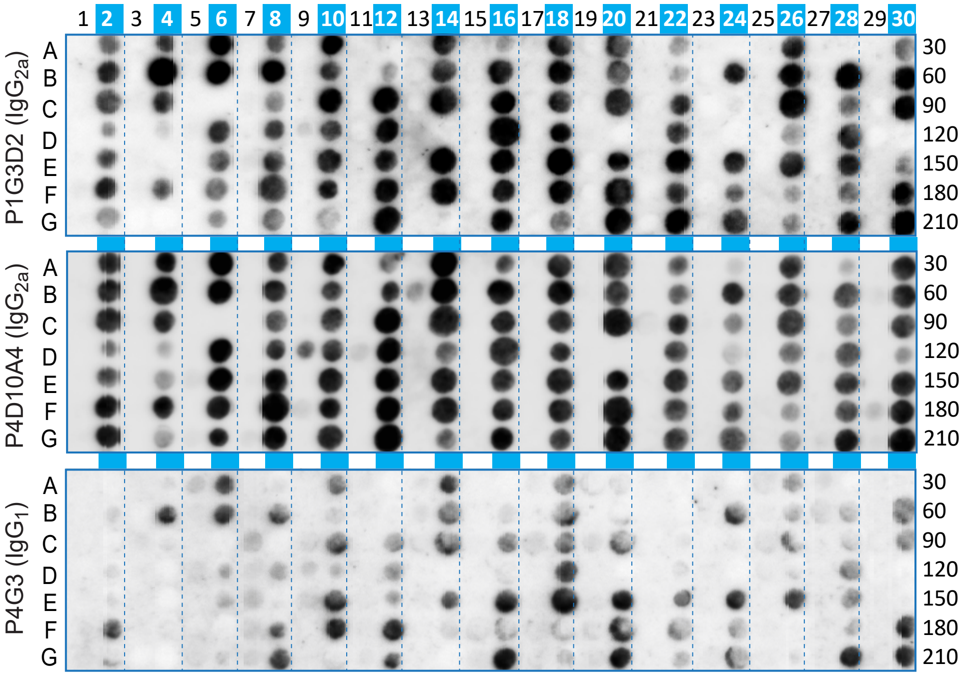


**Supplemental Figure S5. Anti-SDMA hybridoma clones from unimmunized MRL-lpr mice exhibit overlapping and distinct peptide recognition profiles on antigen arrays.**

Three additional hybridoma clones, alongside P4A2F4 and the negative control P2C1C10 (from Figure 4E), were screened against the antigen array. All clones demonstrated exclusive specificity for SDMA-modified peptides (even-numbered columns), with no reactivity against unmodified (odd-numbered columns). Clones P1G3D2 and P4D10A4 (both IgG2a) displayed relatively broad reactivity across the majority of SDMA-containing peptides. In contrast, clone P4G3 (IgG1) recognized a more restricted and distinct subset of SDMA peptides, though with some overlap with the broader clones.

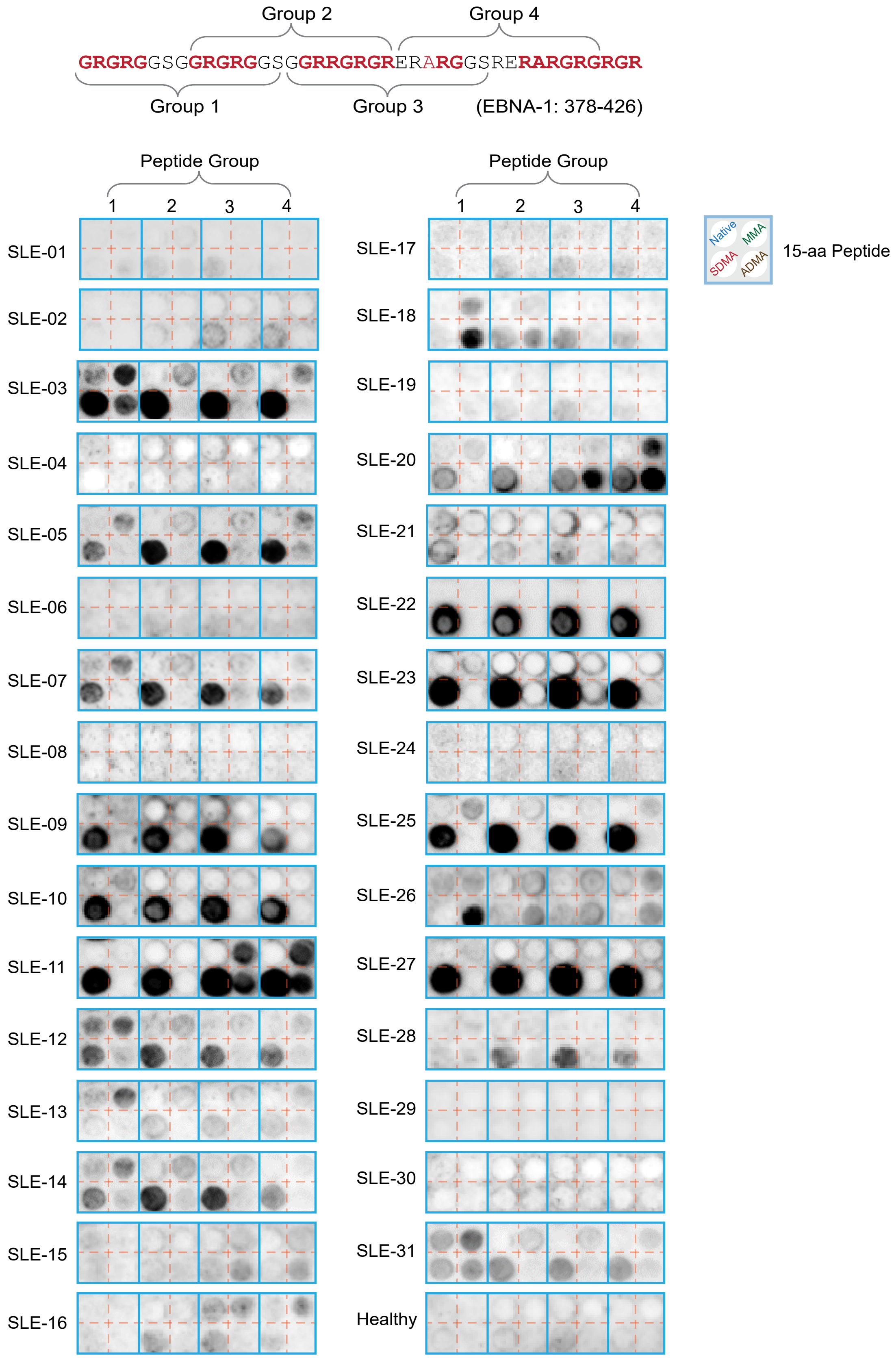


**Supplemental Figure S6. Specific antibody reactivity to SDMA-modified RG repeat epitopes derived from EBNA1.** Four EBNA1-derived RG-repeat peptides, with or without, MMA, SDMA, or ADMA modifications, were probed with individual plasma samples from 31 SLE patients (SLE-01 to SLE-31) from the discovery cohort, following the same protocol as in Figure 2H. Dominant SDMA-specific reactivity was detected in 13 of 31 samples (SLE-03, 05, 07, 09, 10, 11, 12, 14, 20, 22, 23, 25, and 27), whereas healthy control plasma showed no detectable signals, as expected. Signal intensities varied across both patient samples and the four peptides. Notably, weak cross-reactivity with MMA was observed in SLE-03, 11, 12, and 20, and with ADMA in SLE-11, 18, 20, and 26, though these signals were generally lower than those against SDMA.

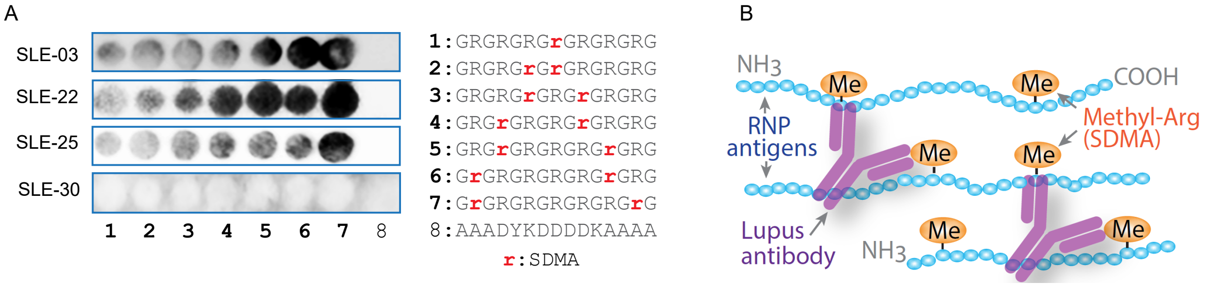


**Supplemental Figure S7. Propensity of anti-SDMA antibodies to form supersized immune complexes.**

A. Effect of SDMA epitope spacing on IgG binding. A series of peptides (#1–#7), each bearing two SDMA residues separated by 1–11 amino acids (right panel), were probed with plasma from SLE-03, SLE-22, SLE-25 and SLE 30. Spot #8 was a negative control peptide with no SDMA modifications. Antibody reactivity increased with greater inter-SDMA distance across all positive samples, albeit to varying degrees, suggesting that close proximity hinders simultaneous binding of two IgG molecules due to steric interference. As expected, SLE-30 plasma showed no no reactivity to any SDMA-containing peptides.

B. Schematic model. Based on the spacing-dependent binding principle, bifunctional IgG molecules that engage multiply methylated lupus antigens may promote the assembly of large immune complexes (supersized immunocomplexes).

**Supplemental Figure S8. Human Smith Antigen Family of snRNPs (RG highlighted).**

```
SmD2        --------------MSLLNKPKSEMTPEELQKREEEEFNTGPLSVLTQSVKNNTQVLINC        46
SmE         MAYRGQGQKVQKVMVQPINLIFR---------------------YLQ--NRSRIQVWLYE        37
SmB/B'      ---------------MTVGKSSK----------------------ML--QHIDYRMRCIL        21
SmN         ---------------MTVGKSSK----------------------ML--QHIDYRMRCIL        21
SmD1        ------------------MKLVR---------------------FLM--KLSHETVTIEL        19
SmD3        ---------------MSIGVPIK---------------------VLH--EAEGHIVTCET        22
SmF         --------------MSLPLNPKP---------------------FLN--GLTGKPVMVKL        23
SmG         ----------------MSKAHPP---------------------ELK--KFMDKKLSLKL        21
                                                         :         :

SmD2        RNNKKLLGRVKAFDRHCNMVLENVKEMWTEVPKSGKGKKKSKPVNKDRYISKMFLRGDSV        106
SmE         QVNMRIEGCIIGFDEYMNLVLDDAEEIHSK---------T----KSRKQLGRIMLKGDNI        84
SmB/B'      QDGRIFIGTFKAFDKHMNLILCDCDEFRKIKPKNSKQAER----EEKRVLGLVLLRGENL        77
SmN         QDGRIFIGTFKAFDKHMNLILCDCDEFRKIKPKNAKQPER----EEKRVLGLVLLRGENL        77
SmD1        KNGTQVHGTITGVDVSMNTHLKAVKMTLKN--------------REPVQLETLSIRGNNI        65
SmD3        NTGEVYRGKLIEAEDNMNCQMSNITVTYRD--------------GRVAQLEQVYIRGSKI        68
SmF         KWGMEYKGYLVSVDGYMNMQLANTEEYIDG--------------ALSGHLGEVLIRCNNV        69
SmG         NGGRHVQGILRGFDPFMNLVIDECVEMATS--------------GQQNNIGMVVIRGNSI        67
            . .    * .   :   *  :                            :  : :: ..:

SmD2        IVVLRNPLIAGK------------------------------------------------        118
SmE         TLLQSVSN----------------------------------------------------        92
SmB/B'      VSMTVEGPPPKDTGIARVP----------LAGAAGG-----PGIGRAAGRGIPAGVPMPQ        122
SmN         VSMTVEGPPPKDTGIARVP----------LAGAAGG-----PGVGRAAGRGVPAGVPIPQ        122
SmD1        RYFILPDSLPLDTLLVDVEPKVKSKKREAVAGRGRG-----RGRGRGRGRGRGRGGPRR-        119
SmD3        RFLILPDMLKNAPMLKSMKNKNQG----SGAGRGKAAILKAQVAARGRGRGMGRGNIFQK        124
SmF         LYIRGVEEEEEDGEMRE-------------------------------------------        86
SmG         IMLEALERV---------------------------------------------------        76
              .

SmD2        ------------------------------------------------------------        118
SmE         ------------------------------------------------------------        92
SmB/B'      APAGLAGPVRGVGGPSQQVMTPQGRGTVAAAAAAATASIAGAPTQYPPGRGGPPPPMGRG        182
SmN         APAGLAGPVRGVGGPSQQVMTPQGRGTVAAAAVAATASIAGAPTQYPPGRGTPPPPVGRA        182
SmD1        ------------------------------------------------------------        119
SmD3        RR----------------------------------------------------------        126
SmF         ------------------------------------------------------------        86
SmG         ------------------------------------------------------------        76


SmD2        ----------------------------------------------------------          118
SmE         ----------------------------------------------------------          92
SmB/B'      APPPGMMGPPPGMRPPMGPPMGIPPGRGTPMGMPPPGMRPPPPGMRGPPPPGMRPPRP          240
SmN         TPPPGIMAPPPGMRPPMGPPIGLPPARGTPIGMPPPGMRPPPPGIRGPPPPGMRPPRP          240
SmD1        ----------------------------------------------------------          119
SmD3        ----------------------------------------------------------          126
SmF         ----------------------------------------------------------          86
SmG         ----------------------------------------------------------          76
```

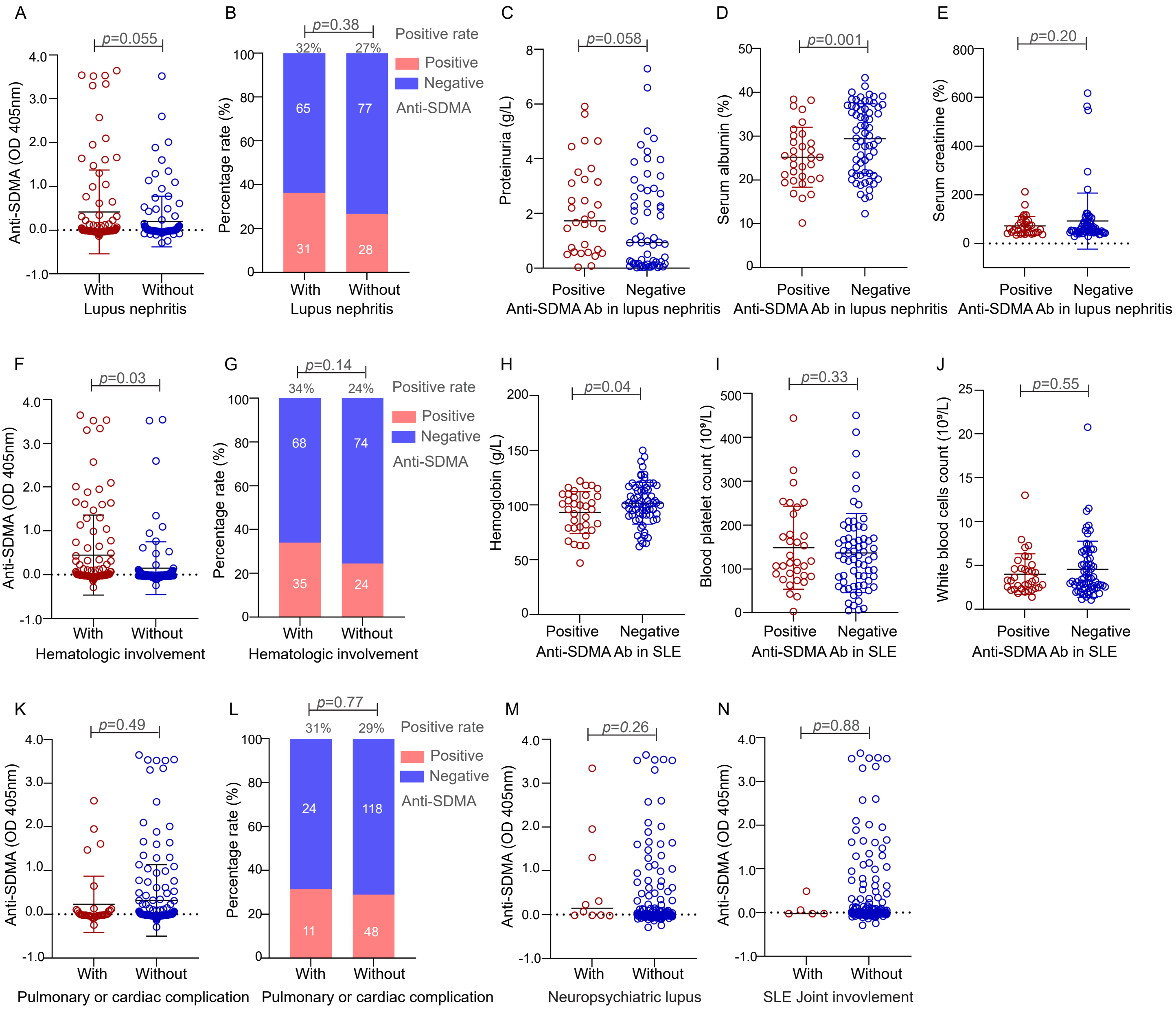


**Supplemental Figure S9. Anti-SDMA seropositivity in correlation with SLE organ involvement.**
The validation cohort of 201 SLE subjects were investigated for the correlation of anti-SDMA seropositivity and/or titer with kidney (A-E), hematologic (F-J), pulmonary or cardiac (K-L), neuropsychiatric (M), and joint involvements (N).
